# Liver Disease Reveals KIF12 as a Critical Regulator of Mitochondria, Lysosome and Cilia Localization

**DOI:** 10.64898/2026.02.24.707575

**Authors:** Anubha Seth, Joseph Brancale, Nina Dashti-Gibson, Rodrigo M. Florentino, Lanuza A. P. Faccioli, Zhenghao Liu, Chigoziri Konkwo, Hyunseok Hong, Alejandro Soto-Gutierrez, Silvia Vilarinho

**Affiliations:** Departments of Internal Medicine, Genetics and Pathology, Yale School of Medicine, New Haven, CT; Department of Pathology, Center for Transcriptional Medicine, University of Pittsburgh, Pittsburgh, PA

## Abstract

Pathogenic variants in kinesin family member 12 (KIF12) cause pediatric liver disease, yet the cellular mechanisms underlying this phenotype remain unknown. Here we show that *KIF12* expression in the healthy human liver is primarily detected in biliary epithelial cells, as revealed by single cell RNA-sequencing data. To investigate its role in biliary pathology, we introduced a homozygous KIF12 p.Arg219* mutation into induced pluripotent stem cells (iPSCs), which were differentiated into 2D cholangiocyte-like cells (iCCs) and 3D biliary organoids. Pioneering single-molecule fluorescence microscopy in live iCCs, we observed wildtype KIF12 co-localizing with microtubules, consistent with its predicted role as a microtubule-associated motor protein. Our data reveals that KIF12 dysfunction causes abnormal perinuclear clustering of mitochondria and lysosomes, and mislocalization of primary cilia in cholangiocytes. Restoration of wildtype KIF12 expression in mutant KIF12 iCCs rescued organelle positioning and normalized GGT activity. This study uncovers a novel link between KIF12 dysfunction and organelle dynamics in human cholangiocytes, extending our understanding of kinesin roles beyond their established functions in neuronal systems. Our findings provide new insights into the pathogenesis of KIF12-related cholestatic liver disease and lay the groundwork for developing targeted genetic therapies.

## INTRODUCTION

The liver is the largest solid organ in the human body and comprises a variety of cell types with a diverse array of cellular and molecular functions that are essential for survival. While hepatocytes account for the majority of liver cells, biliary cells, also known as cholangiocytes, line the intrahepatic and extrahepatic bile ducts. Cholangiocytes are involved in the transport and modification of the primary bile produced by the hepatocytes. Cholestasis results from impaired bile drainage, leading to stasis and retention of hepatic byproducts in the blood circulation that are normally excreted into bile, resulting in liver injury.

While rare in isolation, as a group, genetic cholestatic liver diseases(1) pose a significant disease burden due to incapacitating symptoms of chronic cholestasis, progression to end-stage liver disease, and the need for life-saving liver transplantation. Cholestatic liver disease is a leading cause for pediatric liver transplant largely due to limited effective therapeutic options. Hence, the discovery of, and investigations into the pathobiology of distinct forms of cholestatic liver disease represent not only an unmet medical need but also an opportunity to advance our understanding of liver biology. We and others have discovered that rare recessive mutations in *KIF12* cause high GGT cholestasis during childhood(2, 3). Since then, twenty-seven individuals with KIF12-mediated cholestatic liver disease have been reported(2-12). However, the underlying mechanism(s) remain elusive, and the available mouse models are limited as they do not recapitulate the high GGT cholestasis phenotype seen in humans(13, 14). KIF12 encodes the kinesin family member 12, one of over 40 human KIF genes, and is predicted to function as a microtubule-associated protein. Despite nearly four decades of research since the first kinesin was discovered, the role of kinesins in liver disease remains mostly unexplored and the function of KIF12 in biliary cells is unknown.

In this study, we generated cholangiocyte-like cells (iCCs) and a biliary organoid model derived from human induced pluripotent stem cells (hiPSC) to investigate human KIF12 dysfunction and advance our understanding of the pathogenesis of this novel and poorly understood genetic cholestatic liver disease.

This study uncovers the role of KIF12 in the regulation of organelle dynamics in human cholangiocytes, offering new insights into the fundamental mechanisms governing subcellular organelle organization. This study lays the groundwork for future research into the contribution of organelle mislocalization to cholestatic liver disease and holds promise for developing targeted therapeutic interventions aimed at halting disease progression and potentially averting the need for liver transplantation in these patients.

## RESULTS

### KIF12 is selectively expressed in cholangiocytes in the human liver

Given that human KIF12-related disease causes high GGT cholestasis, we investigated which cells within the liver express *KIF12*. For this purpose, we took advantage of a human liver single cell atlas(15), which is comprised of over 36,000 cells from 28 non-diseased livers. We found that *KIF12* is selectively expressed in cholangiocytes within the healthy human liver (Figure 1A). Additionally, analysis of publicly available single cell RNA-sequencing (scRNA-seq) from primary human intrahepatic, extrahepatic, and gallbladder cholangiocytes validated *KIF12* expression in these tissues (Figure 1A)(16). While *KIF12* is one of the most abundantly expressed KIFs in human cholangiocytes, these cells also express *KIF5B*, *KIFC3*, *KIF1C*, *KIF3B*, among other KIFs expressed at a lower level (Figure S1).

**Figure 1.**
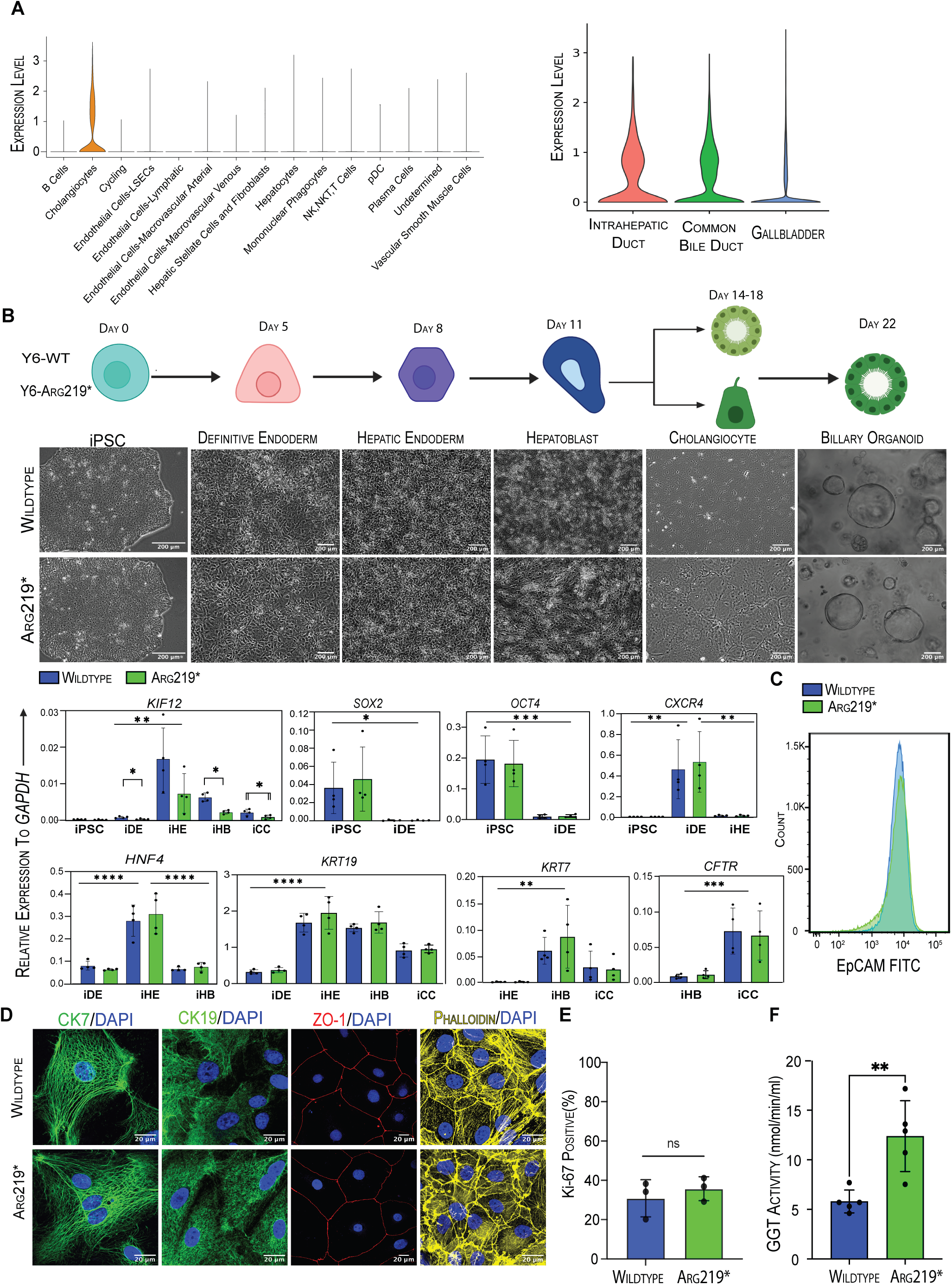
Human *KIF12*-related biliary disease cell-based model. (A) *KIF12* expression across liver cell types using integrated scRNA-seq data from 28 non-diseased human livers, and in cholangiocytes from human intrahepatic bile duct, common bile duct, or gallbladder using publicly available data(16). (B) Schematic overview of iPSC-derived differentiation into cholangiocyte-like cells with representative images at each stage and mRNA expression of *KIF12* and other markers at key stages of Y6 iPSC biliary specification relative to the housekeeping gene *GAPDH* (n=4). (C) Surface expression of epithelial marker EpCAM in Y6 iCCs on final day of differentiation (n=3). (D) Representative images of CK7 and CK19 (green), ZO-1 (red), and phalloidin (yellow) immunostaining in KIF12 wildtype and Arg219* Y6 iCCs. (E) Percentage of Ki67 positive cells in wildtype and KIF12 Arg219* Y6 iCCs. >2000 nuclei per differentiation (n=3). (F) GGT activity measured in wildtype and KIF12 Arg219* iCCs (n=5). iPSC, induced pluripotent stem cell; iDE, definitive endoderm; iHE, hepatic endoderm; iHB, hepatoblast; iCC, cholangiocyte-like cell. Bar graphs represent mean ± SD. *p ≤ 0.05, **p≤0.01, ***p≤0.001, ****p≤0.0001.

### Y6 iPSC differentiation into 2D cholangiocyte-like cells and 3D biliary organoids reveal KIF12 upregulation at hepatic endoderm stage

Taken advantage of previously described protocols(17-20), we successfully differentiated Y6 fibroblast-derived hiPSC(21) into cholangiocyte-like cells (iCCs) using a sequential differentiation procedure through the following stages: definitive endoderm (iDE), hepatic endoderm (iHE), hepatoblast (iHB), and terminating in iCCs (Figure 1B). Using CRISPR/Cas9 gene editing, we generated a homozygous premature termination mutation in the *KIF12* gene, targeting the 219^th^ amino acid residue (chr9:116857353, C>T, Arg219*) within the Y6 hiPSC line. To date, this pathogenic mutation represents the most reported disease-causing mutation in *KIF12*(2, 4). Both wildtype and KIF12-mutant Arg219* iPSCs similarly differentiated into iCCs (Figure 1B). These stages of differentiation were confirmed by downregulation of *SOX2* and *OCT4* and upregulation of *CXCR4* in iDE, *HNF4A*, *KRT19* in HE, *KRT7* and *CFTR* in iCC (Figure 1B). Importantly, we found that *KIF12* expression is upregulated over twenty-fold from iDE to iHE, and its expression persists through iHB and iCC stages (Figure 1B). Consistent with findings from another independent group(4), *KIF12* mRNA expression is decreased, but still detected in cells harboring the homozygous truncating mutation (p.Arg219*) in *KIF12*, suggesting that it partially escapes mRNA decay. Both wildtype and Arg219* *KIF12* iCCs showed no difference in expression of EPCAM, biliary markers CK7 and CK19, ZO-1, and also maintained normal-appearing F-actin architecture as revealed by phalloidin staining (Figure 1C,D). No significant variance in either the differentiation of iPSCs into iCCs or their rate of proliferation (Figure 1E) with or without p.Arg219* *KIF12* mutation was noted.

GGT activity is elevated approximately two-fold in p.Arg219* iCCs as compared to their wildtype controls (Figure 1F) recapitulating the pattern observed in the human KIF12-related human disease. Furthermore, p.Arg219* and wildtype biliary organoids can form, grow, and express similar levels of mature biliary markers CK7 and CK19 (Figure S2A). Biliary organoids from both genotypes show no apparent difference in swelling upon exposure to secretin (Figure S2B).

To further characterize the role of *KIF12* in our cholangiocyte model, we performed RNA-sequencing of Y6 wildtype and p.Arg219* *KIF12* iHE, the stage where *KIF12* upregulation was detected, and iCCs. Principal component analysis (PCA) shows cells of both genotypes being largely similar at the iHE stage, while they mildly diverge at the iCC stage (Figure S3A). Additionally, PCA showed that Y6 iCCs are more transcriptionally similar to adult human cholangiocytes(22-24) than other cholangiocyte cell models(25, 26)(Figure S3B). Furthermore, gene score analysis confirmed that Y6 wildtype and p.Arg219* *KIF12* iCCs are most similar to intrahepatic cholangiocytes as compared to extra-hepatic cholangiocytes (Figure S3C).

Using statistically significant differentially expressed genes (DEGs) in iCCs, we performed gene ontology enrichment analyses for cellular components, biological processes, and molecular function between Y6 p.Arg219* and wildtype *KIF12* iCCs. We found that DEGs were enriched for genes located within external cell membrane, vesicles, and extracellular matrix; involved in organelle fission and epithelial migration; and functional components of extracellular matrix and enzyme inhibition (Figure S3D-F).

### Tracking movement of wildtype KIF12 in live Y6 iCCs at the single molecule level

We then sought to further characterize the role of KIF12 as a putative microtubule-associated transport protein. Given the lack of a reliable human KIF12 antibody, we developed an overexpression system of wildtype and Arg219* mutant KIF12 in HEK293 cells to assess respective protein expression. As expected, overexpression of p.Arg219* KIF12 showed a very faint band of truncated protein as compared to the full-length wildtype KIF12 (Figure 2A-B). This data supports that p.Arg219* homozygous mutation leads to KIF12 protein degradation and deficiency. We further leveraged our iPSC-based model to pioneering single molecule KIF12 fluorescent microscopy in live iCCs. As the EGFP-overexpression system lacked single molecule resolution, we generated a HaloTag-KIF12 overexpression approach (Figure 2A). We observed that wildtype KIF12 co-localizes strongly with tubulin, which stains for microtubules (Manders’ Correlation of 0.92±0.033, Costes P-Value of 1.00), consistent with its predicted role as a microtubule-associated protein (Figure 2C-D, Figure S4A-B). As expected, HaloTag-Arg219*KIF12 was not detected using single molecule fluorescence microscopy (Figure 2C). Moreover, single molecule kymograph analysis supports an average KIF12 velocity across all molecules around 0.875±0.017 um/s (Figure 2E). In order to further understand KIF12 dynamics, we employed a multi-state modeling designated Spot-ON(27), which suggest that KIF12 might exist in three states: (i) a nonmotile state, (ii) a slow-moving state (0.1um^2^/s) and (iii) a fast state (1.172um^2^/s) (Figure 2F). Following confirmation that KIF12 associates with microtubules in live cholangiocytes, we next examined whether KIF12 dysfunction (p.Arg219*), which prevents KIF12 localization to the cell periphery in iCCs as shown by TIRF microscopy, alters mitochondrial distribution.

**Figure 2.**
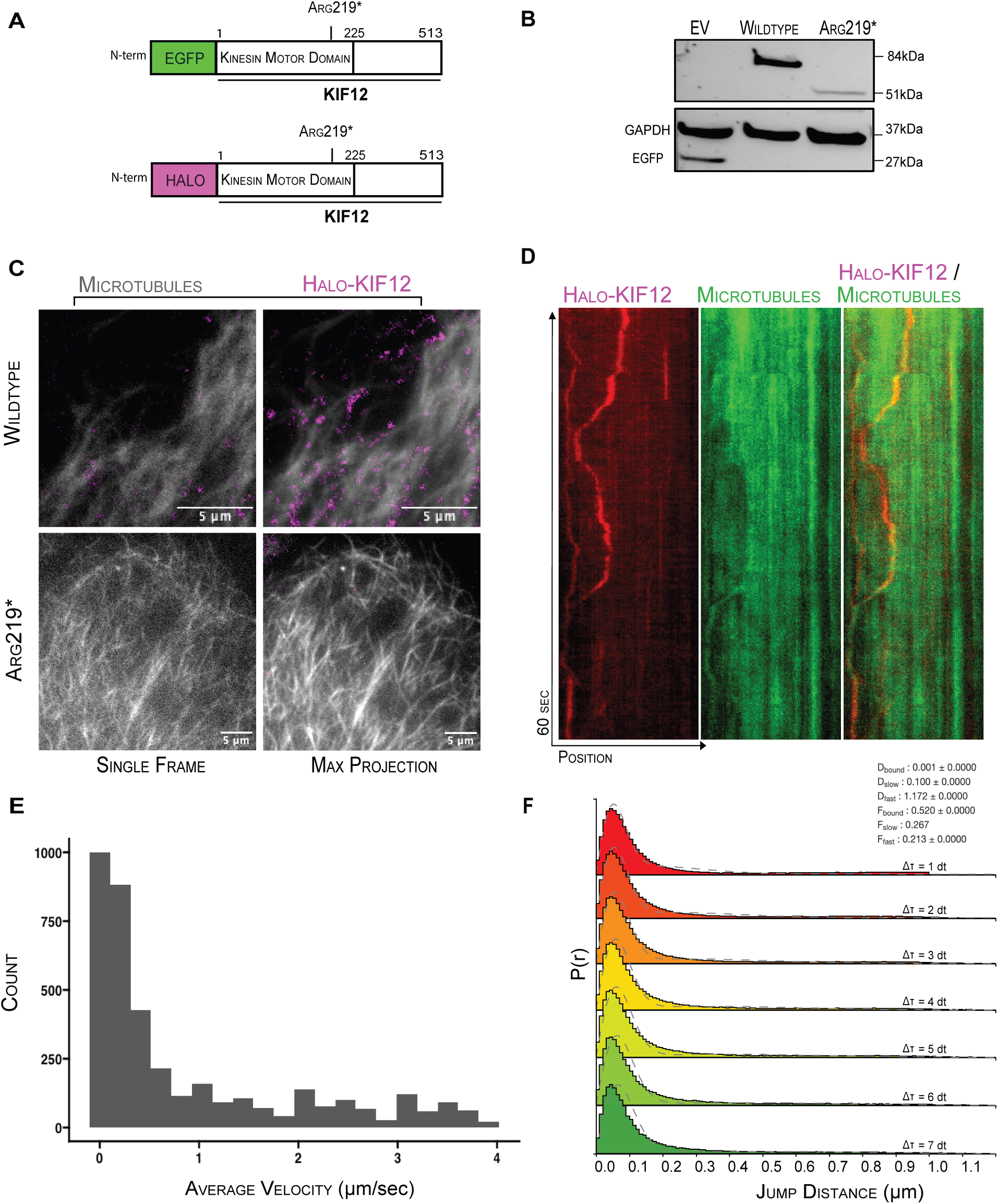
Live microscopy of single KIF12 wildtype molecules in Y6 iCCs. (A) Schematic representation of EGFP or Halo-tagged wildtype and Arg219* constructs. (B) Western blot showing fusion protein expression of EGFP-tagged wildtype and Arg219* KIF12 in HEK293. EV: Empty vector with EGFP tag. (C) Representative images of TIRF microscopy of Tubulin Tracker highlighting microtubules (grey) and wildtype or Arg219* Halo-KIF12 overexpression (magenta) in live iCCs. Both single frames (left panel) and max projection from 1-minute acquisition (right panel) are shown. (D) Representative kymographs of wildtype Halo-KIF12 along a single microtubule. Left kymograph depicts Halo-KIF12, middle kymograph depicts Tubulin Tracker highlighting microtubules, right kymograph is a composite image. (E) Histogram displaying the average velocity (µm/sec) of each count of Halo-KIF12 molecule as obtained by kymograph analysis. n=150 kymographs over 15 images. (F) Spot-ON kinetic modeling of single-molecule Halo-KIF12 tracks are shown (n=15 images). Jump distance (distance traveled between two points in time) distributions modeled between Dt1-7 are shown. Resulting diffusion coefficient and molecular fraction parameters for each state are shown.

### KIF12 orchestrates the spatial organization of mitochondria, lysosomes, and primary cilia in iCCs

Using both cytochrome c immunostaining and overexpression of DsRed containing a mitochondrial localization sequence (mitoDsRed) for endogenous mitochondria labeling, we found that p.Arg219* iCCs show a predominantly perinuclear localization of mitochondria rather than their typical distribution throughout the cytoplasm seen in wildtype iCCs (Figure 3A and Figure S5). This results in a reduction in the mitochondrial footprint in p.Arg219\**KIF12* iCCs as compared to wildtype iCCs (Figure 3A-B and S5). We then sought to investigate whether this mislocalization of mitochondria has an impact in mitochondrial function. For this purpose, we performed Agilent Seahorse XF Pro mitochondrial stress assay, which revealed reduced maximal respiration in p.Arg219* *KIF12* iCCs as compared to wildtype iCCs (Figure 3C,D). These data support that p.Arg219* *KIF12* iCCs exhibit impaired bioenergetic health with decreased mitochondrial function upon exposure to metabolic stress. Given this finding, we generated a second independent iPSC line (FL167) with the same p.Arg219* mutation in *KIF12* (Figure S6A-D) and successfully differentiated into iCCs. These FL167 iCCs validated the abnormal perinuclear mitochondrial location and compromised mitochondria function initially revealed by Y6 line (combined data from Y6 and FL167 lines are shown in Figure 3B-D).

**Figure 3.**
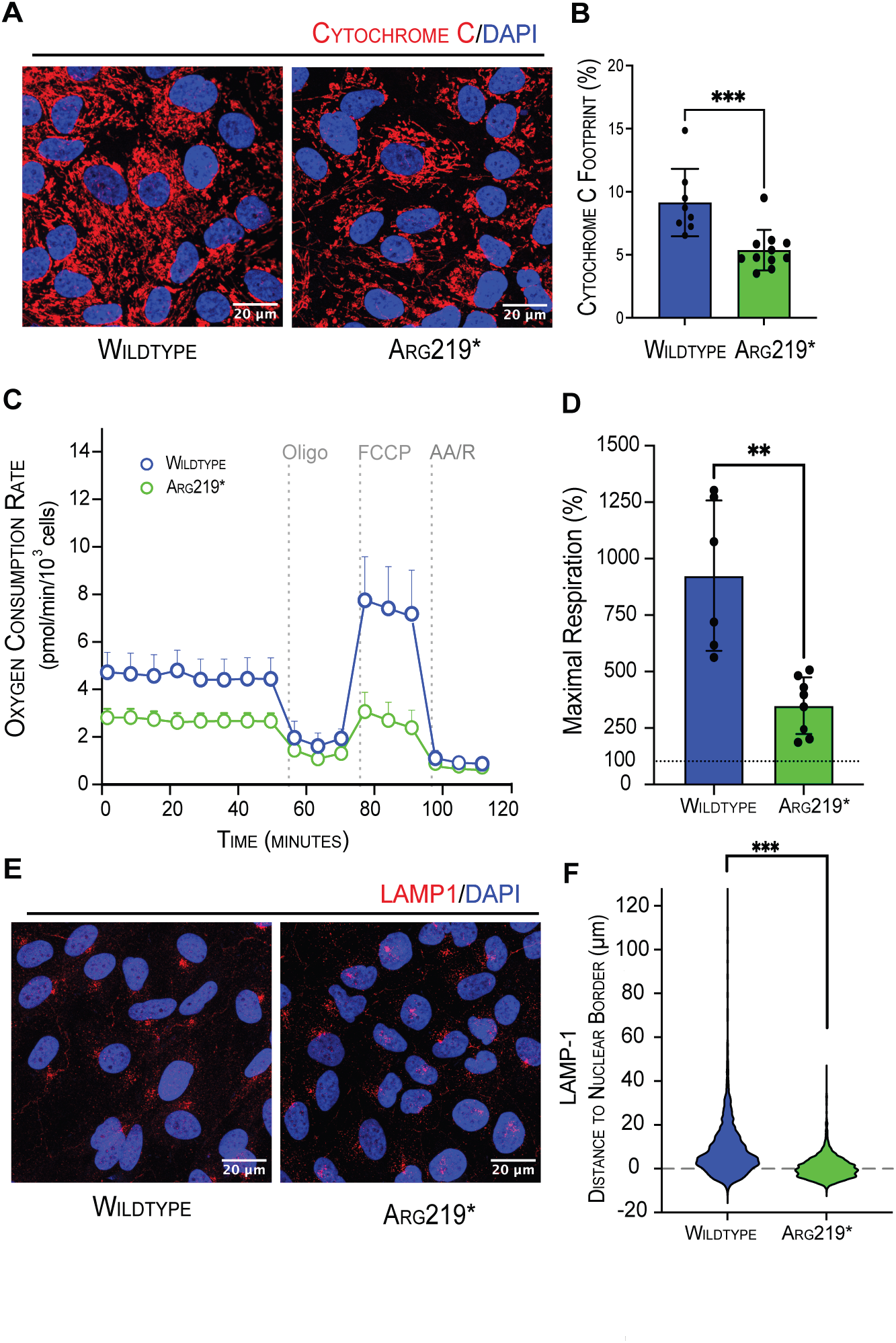
Mitochondrial and lysosome subcellular localization in KIF12 Arg219* iCCs. (A) Representative images of cytochrome c immunostaining showing mitochondrial perinuclear localization and footprint reduction in Arg219* iCCs. (B) Quantification of mitochondrial footprint, with each dot representing the average percentage per experiment. Data are presented as mean ± S.D. for cytochrome C (n≥8). (C) Seahorse mitochondrial stress assay was performed on wildtype and Arg219* iCCs, measuring oxygen consumption rates following sequential injections of the test compounds: oligomycin (Oligo, 1 μM), FCCP (10 μM), and a combination of rotenone (R, 5 μM) with antimycin A (AA, 10 μM). Data are normalized to 1000 cells. Error bars represent S.E.M. (n=6-8). Changes in (D) maximal respiration, 100% represents non-mitochondrial respiration. Error bars indicate mean ± S.D. (E) Representative image of LAMP1 immunostaining in iCCs. (F) Violins represent the distribution of the distance of individual LAMP1^+^ puncta from the nucleus. Quantification includes >2500 LAMP1^+^ puncta per genotype (n=5-7). Error bars indicate mean ± S.D. Significance *p≤0.05, **p≤0.01, ****p≤0.0001.

We next sought to investigate if other organelles had similar localization defects. Employing LAMP1 immunostaining and LAMP1-RFP overexpression for endogenous lysosome labeling, we found that p.Arg219* *KIF12* iCCs also show a predominantly perinuclear localization of lysosomes rather than their conventional distribution throughout the cytoplasm observed in the wildtype iCCs (Figure 3E and Figure S7A). Despite the perinuclear localization, we found no evidence of autophagy dysfunction in Y6 p.Arg219* *KIF12* iCCs relative to respective wildtype iCCs controls (Figure S7B-D). Perinuclear lysosome observation was also validated using Lamp-1 staining in the second independent iPSC line (combined data showed in Figure 3F).

Given that biliary cells are known to have a primary cilium, which is a microtubule-based structure, we proceeded to evaluate cilia in polarized wildtype and p.Arg219* *KIF12* iCCs. While most iCCs from both genotypes show the ability to ciliate (Figure 4A-C), its length appears shorter in p.Arg219* *KIF12* iCCs as compared to wildtype iCCs (Figure 4A-C). Thus, we hypothesize that the trafficking of the cilia might have been compromised in the absence of properly functional KIF12. For that purpose, we employed the IN/OUT assay for the first time to study ciliogenesis in iCCs, with the goal to assess the proportion of intracellular and extracellular cilia(28). This assay revealed that while only a very small proportion of cilia (<3.5%) are intracellular in wildtype iCCs, in p.Arg219* KIF12 iCCs, the proportion of intracellular cilia rises around 4-fold, with approximately 15% of cilia in these cells being intracellular (Figure 4D-E). Additionally, in one of the iPSC lines, cilia appeared mostly bent in Y6 Arg219* iCCs as compared to wildtype Y6 iCCs controls (Figure S8A,B and Videos S1 and S2). Furthermore, we investigated the orientation of cilia in 3D biliary organoids. Whereas in wildtype organoids, cilia were found as expected exclusively on the apical side of wildtype iCCs, in p.Arg219* *KIF12* organoids, cilia were observed on both the apical and basal aspects (Figure 5A and Figure S9A). This observation is also seen in apical-out (inverted) biliary organoids (Figure S9B), with no evidence of cellular polarization defects as supported by normal phalloidin, b-catenin, and CFTR immunostaining (Figure 5A,B and Figure S9A,B). These findings were recapitulated in a second independent FL167 iPSC line (Figure S10A,B).

**Figure 4.**
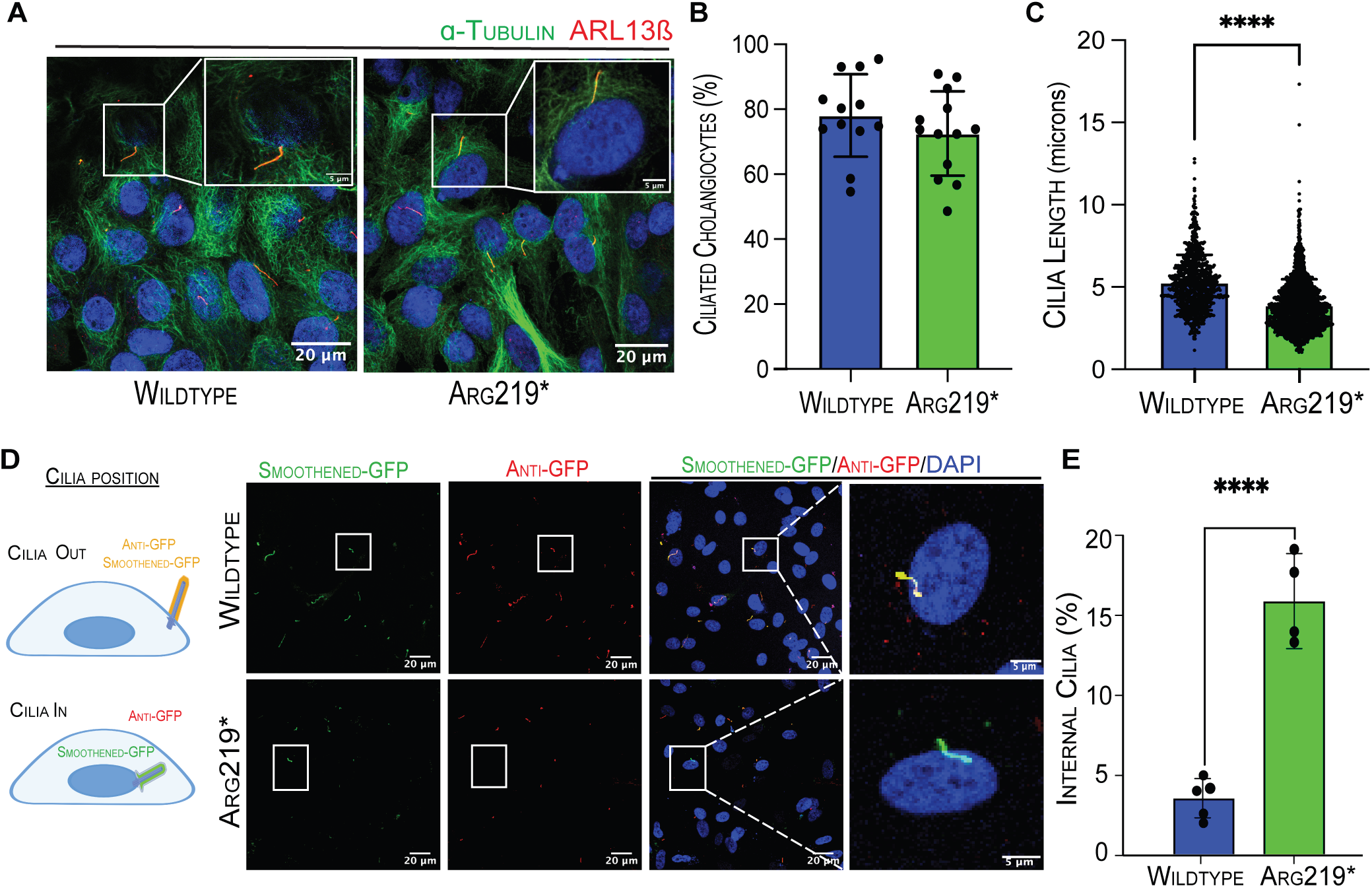
Characterization of primary cilia in *KIF12* Arg219* 2D iCCs. (A) Representative images showing co-immunostaining of cilia in polarized iCCs, with acetylated α-tubulin and ARL13β. Insets show higher magnification images of the cilia region. (B) Quantification of percentage of ciliated iCCs and (C) length of cilia using ARL13β immunostaining with >300 cilia per genotype, (n≥10). (D) Schematic overview of IN/OUT assay data interpretation. Representative images showing overexpression of the ciliary membrane marker Smoothened fused with GFP in iCCs, stained with anti-GFP (red). Extracellular cilia (OUT) are indicated by the co-localization of green and red signals (appearing as yellow), while intracellular cilia (IN) are identified as green given the absence of the anti-GFP signal (red). (E) Quantification of intracellular cilia in wildtype and Arg219* iCCs (n≥4). **p≤0.01, ***p≤0.001,****p≤0.0001.

**Figure 5.**
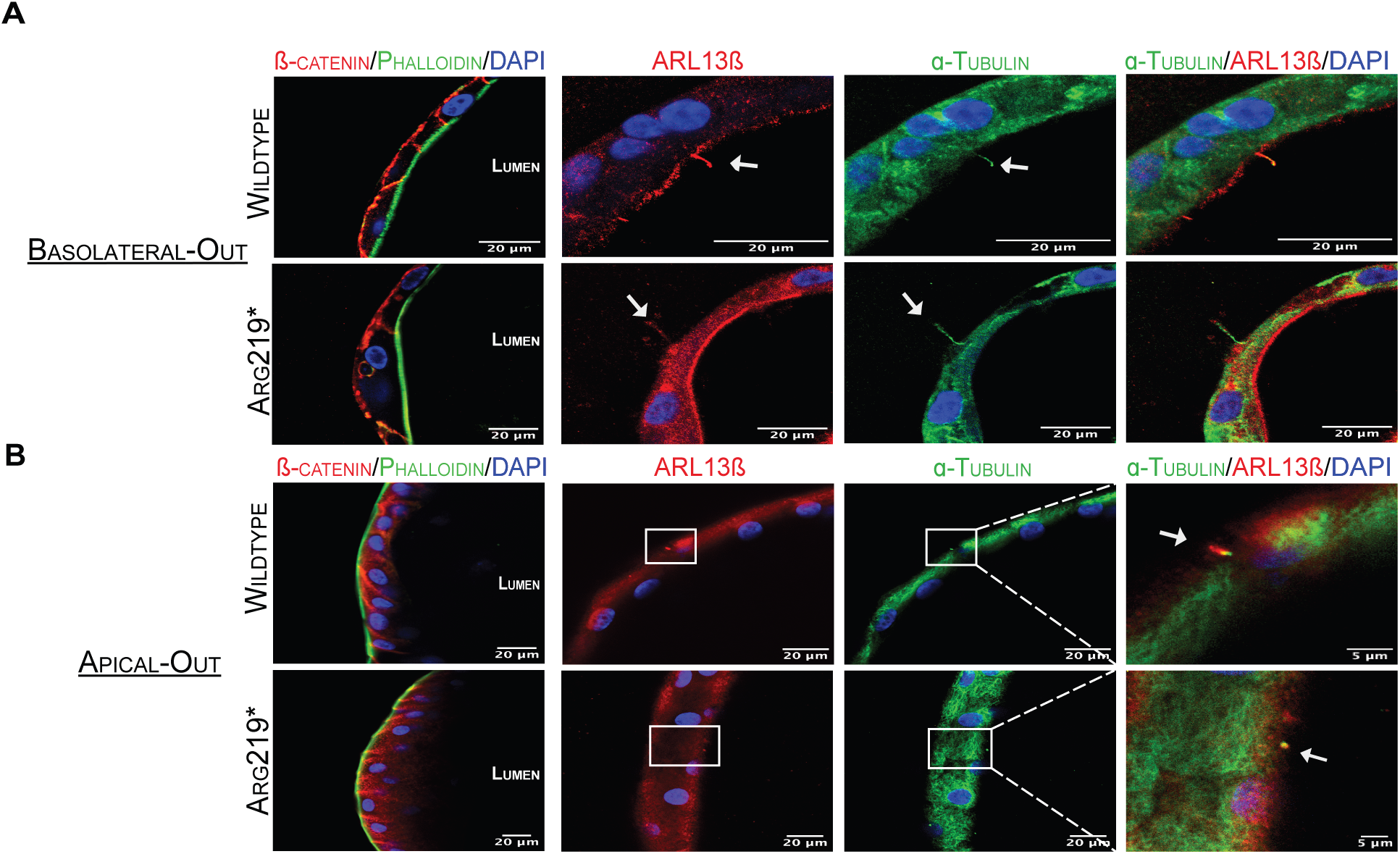
Characterization of primary cilia in *KIF12* Arg219* 3D biliary organoids. (A) Biliary organoids stained with phalloidin (green) and β-catenin (red) illustrate intact apical-basal polarity in both WT and Arg219* iCCs. Co-immunostaining with ciliary markers acetylated α-tubulin (green) and ARL13β (red) reveals apical cilia in wildtype biliary organoids and cilia in apical and basal aspects in KIF12 mutant biliary organoids. (B) ‘Apical-out’ (inverted) biliary organoids showing reversal of phalloidin (green) and β-catenin (red) staining, along with acetylated α-tubulin (green) and ARL13β (red) co-localization on both the apical and basal sides, highlighting cilia misorientation in Arg219* despite maintaining normal apical-basal polarity. White arrows and boxes indicate cilia.

### Exogenous KIF12 rescues organelle mislocalization in Y6 p.Arg219* KIF12 iCCs

Next, we assessed whether overexpression of wildtype KIF12 in Y6 Arg219* mutant iCCs could rescue the organelle mislocalization seen in these cells. Using this overexpression system, we observed a rescue of perinuclear localization of mitochondria and lysosomes (Figure 6A-C). Moreover, overexpression of wildtype KIF12 in p.Arg219* KIF12 iCCs led to recovery of mitochondrial function, as observed by the restoration of maximal respiration measured using mitochondrial stress assay (Figure 6D-E), and normalization of GGT activity (Figure 6F). We also detected restoration of cilia position when wildtype KIF12 is present comparable to the ones seen in wildtype iCCs (Figure S11A,B, and Videos S3 and S4).

**Figure 6.**
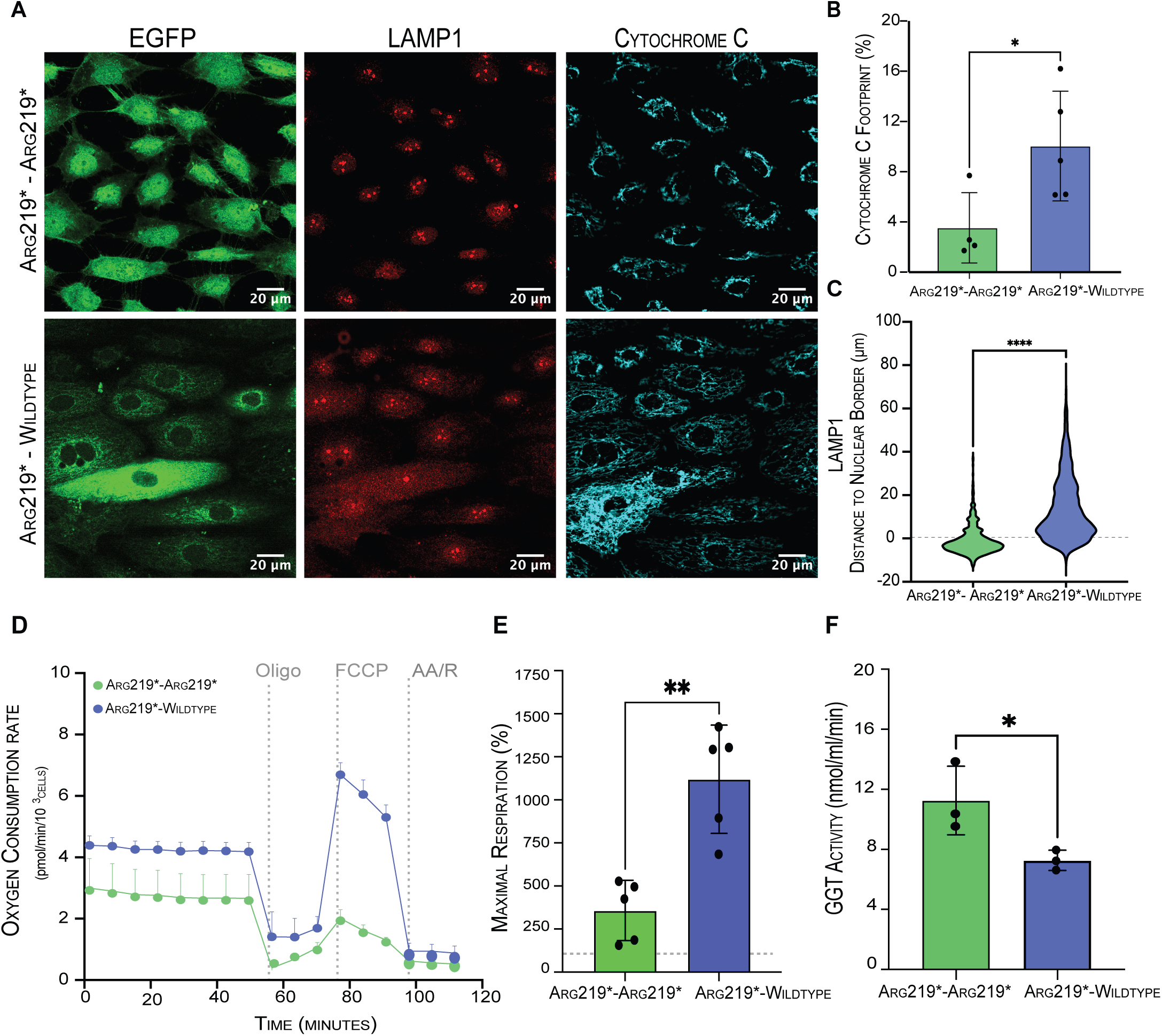
Wildtype KIF12 overexpression rescues organelle mislocalization in Y6 Arg219* iCCs. (A) Representative confocal images of overexpression (OE) of EGFP-tagged Arg219* and Wildtype KIF12 constructs in iCCs co-immunostained with cytochrome C (cyan) and LAMP1 (red) showing normalization of organelle distribution throughout the cytoplasm in Arg219*-Wildtype OE iCCs. (B) Quantification of mitochondrial footprint percentage in Arg219*-Wildtype OE iCCs vs. Arg219*-Arg219* OE (n=5) (C) Distribution of the distance of the individual LAMP1^+^ puncta to the nucleus (quantification includes >750 LAMP1^+^ puncta per genotype, n≥4). Error bars represent mean ± S.D. (D) Mitochondrial stress assay performed on Arg219* iCCs upon overexpression of Arg219*-EGFP or wildtype-EGFP KIF12. Overexpression of wildtype KIF12 restores maximal mitochondrial respiration capacity. Data normalized to 1000 cells; error bars represent SEM. (E) Changes in maximal respiration (100% represents non-mitochondrial respiration) are shown. Error bars indicate mean ± S.D (n=5). (F) GGT activity upon overexpression of wildtype-EGFP or Arg219*-EGFP KIF12 is depicted (n=3). Significance *p≤0.05, **p≤0.01, ***p ≤0.001,****p≤0.0001.

## DISCUSSION

This study reveals KIF12 as a key regulator of organelle position, specifically mitochondria, lysosomes, and cilia, within human biliary cells. Using live-cell imaging, we confirmed that KIF12 co-localizes with microtubules in iCCs. Interestingly, loss of KIF12 function leads to abnormal perinuclear clustering of mitochondria and lysosomes(29), resembling the phenotype previously reported for Kif5b deficiency. Our findings align with the prediction that KIF12 functions as a molecular motor protein associated with microtubules, likely involved in intracellular organelles’ transport. While characterizing the KIF12 we demonstrated its average velocity of 0.875±0.017 um/s and a separate fast-state of 1.172um^2^/s. These speeds are in line with measurements of other kinesins that are similarly involved in cellular cargo transport(30, 31). While p.Arg219* iPSCs differentiate into iCCs and proliferate similarly to wildtype iPSCs and iCCs, KIF12 mutant iCCs displayed a decrease in mitochondrial maximal respiratory capacity and elevated GGT activity, supporting increased oxidative stress(13). These features suggest that KIF12 dysfunction disrupts both the localization and optimal function of key organelles in biliary cells, potentially driving and/or contributing to biliary injury seen in affected patients.

In addition to lysosomal and mitochondrial mislocalization, KIF12 mutant iCCs had prominent ciliary defects. While KIF12 mutant iCCs can form cilia, KIF12 appears crucial for positioning these cilia exclusively on the apical surface of iCCs and ensuring their length and upright orientation. In our model, KIF12 mutant iCCs demonstrate approximately a 4-fold increase in the number of intracellular cilia observed when compared to homeostatic conditions, in addition to a decrease in cilia length.

Primary cilia have been described to play vital roles in cholangiocytes, projecting into the bile duct lumen serving as sensors for changes in the composition, osmolarity and flow of bile(32). Therefore, it is possible that the combination of decreased number of extracellular cilia and the disruption in their length, position and orientation on the apical surface of cholangiocytes contributes to the development of biliary injury and cholestasis observed in human KIF12-related liver disease.

Consistent with our observations, liver tissue from an affected patient revealed absence of primary cilia on cholangiocytes by transmission electron microscopy(7). Similarly, studies on syndromic and non-syndromic cases of biliary atresia - the leading cause of pediatric liver transplantation worldwide - have revealed a decreased number and morphological abnormalities of cholangiocyte primary cilia, with biliary organoids derived from biopsies of patients with biliary atresia demonstrating misorientation of cilia(33-36). These parallels underscore the critical role of primary cilia dysfunction in the pathogenesis of various cholestatic liver disorders and support the addition of KIF12-related liver disease to this group of disorders.

The kinesin superfamily was discovered approximately four decades ago and has primarily been studied in neurons due to their lengthy and asymmetric structure, necessitating the transport of cargo over considerable distances within the cytoplasm. This study marks the initial comprehensive exploration of one such KIF gene, KIF12, within human biliary cells. By delving into its mechanisms, we establish a crucial connection between the positioning of various organelles, shedding light on their role in liver disease. Furthermore, restoration of wildtype KIF12 expression in mutant KIF12 iCCs rescued organelle positioning and normalized mitochondrial maximal respiratory capacity and GGT activity. These findings demonstrate that the phenotype is reversible, even when wildtype KIF12 is introduced after the defects have already emerged, highlighting the potential for developing targeted genetic therapies for KIF12-related liver disease(37).

## Supporting information

file:///Users/silvlarnho/Desktop/Vilarinho/Manuscripts/KIF12-ms-II/bioRxiv/Supplementary-Information-2-24-2026

file:///Users/silvlarnho/Desktop/Vilarinho/Manuscripts/KIF12-ms-II/bioRxiv/Supplementary-Figures/PDFs/Supplementary-Figure-1.pdf

file:///Users/silvlarnho/Desktop/Vilarinho/Manuscripts/KIF12-ms-II/bioRxiv/Supplementary-Figures/PDFs/Supplementary-Figure-2.pdf

file:///Users/silvlarnho/Desktop/Vilarinho/Manuscripts/KIF12-ms-II/bioRxiv/Supplementary-Figures/PDFs/Supplementary-Figure-3.pdf

file:///Users/silvlarnho/Desktop/Vilarinho/Manuscripts/KIF12-ms-II/bioRxiv/Supplementary-Figures/PDFs/Supplementary-Figure-4.pdf

file:///Users/silvlarnho/Desktop/Vilarinho/Manuscripts/KIF12-ms-II/bioRxiv/Supplementary-Figures/PDFs/Supplementary-Figure-5.pdf

file:///Users/silvlarnho/Desktop/Vilarinho/Manuscripts/KIF12-ms-II/bioRxiv/Supplementary-Figures/PDFs/Supplementary-Figure-6.pdf

file:///Users/silvlarnho/Desktop/Vilarinho/Manuscripts/KIF12-ms-II/bioRxiv/Supplementary-Figures/PDFs/Supplementary-Figure-7.pdf

file:///Users/silvlarnho/Desktop/Vilarinho/Manuscripts/KIF12-ms-II/bioRxiv/Supplementary-Figures/PDFs/Supplementary-Figure-8.pdf

file:///Users/silvlarnho/Desktop/Vilarinho/Manuscripts/KIF12-ms-II/bioRxiv/Supplementary-Figures/PDFs/Supplementary-Figure-9.pdf

file:///Users/silvlarnho/Desktop/Vilarinho/Manuscripts/KIF12-ms-II/bioRxiv/Supplementary-Figures/PDFs/Supplementary-Figure-10.pdf

file:///Users/silvlarnho/Desktop/Vilarinho/Manuscripts/KIF12-ms-II/bioRxiv/Supplementary-Figures/PDFs/Supplementary-Figure-11.pdf

file:///Users/silvlarnho/Desktop/Vilarinho/Manuscripts/KIF12-ms-II/bioRxiv/Movies/Supplementary-Movie-S1.mov

file:///Users/silvlarnho/Desktop/Vilarinho/Manuscripts/KIF12-ms-II/bioRxiv/Movies/Supplementary-Movie-S2.mov

file:///Users/silvlarnho/Desktop/Vilarinho/Manuscripts/KIF12-ms-II/bioRxiv/Movies/Supplementary-Movie-S3.mov

file:///Users/silvlarnho/Desktop/Vilarinho/Manuscripts/KIF12-ms-II/bioRxiv/Movies/Supplementary-Movie-S4.mov

## ACKNOWLEDGMENTS

We thank Mr. Jason Thomson and Dr. Caihong Qiu from the Yale Stem Cell Center for the Y6 iPSC line and technical advice; Dr. Mariangela Amenduni, Dr. Romina Fiorotto and Dr. Carol Soroka from the Yale Liver Center for technical advice; Dr. Felix Rivera-Molina and Dr. Derek Toomre from the Yale CINEMA Lab for both the IN/OUT lentiviral plasmid as well as for technical advice on TIRF microscopy; Mrs. Rebecca Cardone from Yale Chemical Metabolism Core for technical support with Seahorse XF Pro Analyzer; Ms. Morven Graham and Dr. Xinran Liu from the Yale CCMI electron microscopy core for technical assistance; Ms. Teresa Brevini and Dr. Fotios Sampaziotis for facilitating access to previously published scRNA-Seq. S.V. is supported by Doris Duke Charitable Foundation Grant #2019081 and the NIH/NIDDK (R01 DK131033). This work was also supported in part by the Yale Liver Center (P30DK034989). This work was supported by NIH grant DK099257 to A.S.G. and internal funds from the Center for Transcriptional Medicine at the University of Pittsburgh.

## AUTHOR CONTRIBUTIONS

S.V. conceptualized, designed, supervised, and obtained funding to conduct the study. A.S.G. supervised and funded a part of this study. A.S., J.B., N.D-G, R.M.F., L.A.P.F., Z.L. and H.H. designed and conducted experiments, and performed respective analysis. J.B. and C.K. performed RNA sequencing analysis, with input from all authors. A.S, J.B., N.D-G, C.K. S.V. contributed to the writing of the first draft, and all authors approved the final version.

## DATA AND CELL AVAILABILITY

RNA-sequencing data have been deposited in the National Center for Biotechnology Information’s Gene Expression Omnibus (accession no. GSE278954). Materials generated in this study are available upon reasonable request and completion of a materials transfer agreement with Yale University or University of Pittsburgh.

## MATERIALS AND METHODS

### Generation and maintenance of iPSC lines

The Y6 human female iPSC line was obtained from the Yale Stem Cell Center and previously reported by Dash et al.(21). CRISPR/Cas9 technology was used to introduce a C>T nucleotide substitution at chr9:116857353 leading to a premature termination of KIF12 (p.Arg219*). Two guide RNA sequences were used to induce DNA break; guide #1: TGCGATATGCAAGCCGAGCT, guide #2: GCGATATGCAAGCCGAGCTC. An additional synonymous C>A nucleotide substitution at chr9:116857349 was used to prevent re-cutting of target region after gene editing. Two independent homozygous p.R219* clones (#13 and #42) were generated and confirmed by Sanger sequencing. The FL167 human male iPSC line was obtained from the Center for Transcriptional Medicine at University of Pittsburgh and used to introduce the same mutation reported above (p.Arg219*). iPSCs were cultured in feeder-free conditions on Matrigel (Corning) or Geltrex (Gibco) coated plates with mTESR Plus media (STEMCELL Technologies) and passaged using Dispase in DMEM/F12 (STEMCELL Technologies).

### Differentiation of cholangiocyte-like cells from iPSCs

The differentiation of iCCs from iPSCs was carried out according to described protocols(17-20) with modifications. iPSCs were dissociated into single-cell suspension using Accutase (STEMCELL Technologies), seeded on Matrigel-coated 6-well plates at a density of 8x10^5^ cells/well, and cultured in mTESR Plus with 10 μM Y-27632 (STEMCELL Technologies) for 24 hours. To induce definitive endoderm specification, cells were cultured using the STEMdiff Definitive Endoderm Kit (STEMCELL Technologies) for four days according to the manufacturer’s protocol. On day 5, media was changed to KnockOut DMEM (Gibco) with 20% KnockOut Serum Replacement (Gibco), 1% DMSO (Sigma), 1% Non-Essential Amino Acids (Gibco), 1 mM L-glutamine (Gibco), 0.1 mM ß-mercaptoethanol (Gibco), and 1% Pen-Strep (Gibco) for three days to induce hepatic endoderm specification. On day 8, media was changed to RPMI (Gibco) with 2% B27 Supplement (Gibco), 1% Non-Essential Amino Acids, 25 mM HEPES (Gibco), 1% Pen-Strep, 30 ng/mL FGF4 (R&D Systems), 50 ng/mL EGF (Peprotech), 25 ng/mL HGF (Peprotech), and 0.1 μM Retinoic Acid (Sigma) for three days to generate hepatoblasts. On day 11, hepatoblasts were detached using Accutase, split 1:3 onto Collagen I-coated plates (STEMCELL Technologies), and cultured in plating media: 1:1 William’s E/Ham’s F12 (Gibco) with 10% FBS-heat inactivated (Gibco), 0.1% BSA-fatty acid free (Sigma), 1 mM L-glutamine, and 1% Pen-Strep for four hours to promote cell attachment. After four hours, media was changed to biliary differentiation (BD) media, which consisted of William’s E/Ham’s F12 with 0.1% BSA, 0.6 mM Vitamin C (DOT Scientific), 1% Insulin-Transferrin-Selenium (Gibco), 1 mM Sodium Pyruvate (Gibco), 50 nM Triiodo-L-Thyronin (Sigma), 1 mM L-glutamine, and 1% Pen-Strep. After 24 hours, BD media was supplemented with 25 ng/mL EGF and 50 ng/mL HGH (Peprotech) for three days. On day 15, media was replaced with expansion media (EM), consisting of Advanced DMEM/F12 (Gibco) with 10% Rspondin-1 conditioned media, 2% B27 Supplement, 1% GlutaMAX (Gibco), 1% Pen-Strep, 1% N-2 Supplement (Gibco), 10 mM Nicotinamide (Sigma), 1 mM N-acetylcysteine (Sigma), 10 μM Forskolin (Sigma), 5 μM A83-01 (Sigma), 10 nM Gastrin (Sigma), 100 ng/mL FGF10 (Peprotech), 50 ng/mL EGF, and 25 ng/mL HGF for three days to complete biliary differentiation. iPSC-derived iCCs were maintained in culture for up to 15 passages on Collagen I-coated plates in EM as described above. Cells were passaged using 2 mg/mL Collagenase Type 2 (Worthington Biochemical).

### Generation of organoids from iPSCs

On day 11 of the differentiation, hepatoblasts were plated in 3D culture to generate biliary organoids in parallel to cholangiocyte monolayers. After detaching hepatoblasts with Accutase, cells were suspended in a mixture of 75% Matrigel 25% BD media as described above at a density of 6x10^5^ cells/mL. Droplets of 50 μL were plated in the center of each well, allowed to solidify at 37 °C for 30 min, and overlaid with BD media. After 24 hours, media was supplemented with 50 ng/mL HGH and 25 ng/mL EGF for three days, followed by 10 ng/mL IL-6 (R&D Systems) and 25 ng/mL EGF for three days, generating cholangiocyte-like cells (iCCs). On day 18, the media was switched to EM as previously described. Organoids were also generated directly from iCCs, which were plated in a mixture of 75% Matrigel and 25% Wnt3a-conditioned media and maintained for 24 hours. After this period, the organoids were transferred to EM. For passaging, the organoids were dissociated mechanically after approximately five days and replated in EM supplemented with Wnt3a-conditioned media as described above, allowing the culture to be sustained for up to 10 passages.

### Apical-out organoids

Apical-out organoids were generated using a previously described protocol(38) with modifications based on the more fragile nature of biliary organoids. After adding cold 5 mM EDTA (Invitrogen) to each well of organoids, contents were transferred to a 5 mL Eppendorf tube and pipetted gently to break up the droplet. Tubes were rocked at 4°C for 45 min to further dissolve the Matrigel and centrifuged at 100xg for 1 min. Organoids were washed twice with Advanced DMEM/F12 and transferred to a 96-well PrimeSurface Ultra-Low Attachment plate (S-BIO) at a density of 1-5 organoids/well or 24 well Ultra-Low Attachment plate (Corning, #3473) at a density of 10-30 organoids/well. Organoids were cultured in EM for at least four days to allow time for polarity inversion and cilia growth.

### Real-time qPCR

RNA was isolated from cells at each stage of differentiation using the RNeasy Plus Mini Kit (Qiagen), and cDNA synthesis was performed using the SuperScript III Reverse Transcriptase Kit (Invitrogen) with Oligo(dT)_12-18_ Primer (Invitrogen), according to the manufacturers’ protocols. Real-time quantitative PCR was performed using TaqMan Fast Advanced Master Mix (Applied Biosystems) and Taqman Gene Expression Assays (Applied Biosystems). Expression levels were normalized to *GAPDH*.

### Immunofluorescence staining and confocal microscopy

For immunofluorescence staining, iCCs were grown on either Collagen I-coated German Coverslips (Electron Microscopy Sciences) or 50 μg/ml Collagen I (Sigma) coated 0.4 μm PET Transwell Inserts (Falcon) until a confluent monolayer had formed. Cells were fixed in either cold 1:1 Methanol/Acetone (Sigma) for 5 min or 4% Paraformaldehyde (Electron Microscopy Sciences) for 20 min, PBS washes (3 x 5 min), blocked and permeabilized in 3% BSA (Sigma) 0.2% Triton X-100 (Thermo Scientific) for 1 hour at RT, and incubated in primary antibody (1:100 in blocking) overnight at 4°C. The next day, cells were washed with PBS, incubated in secondary antibody (1:500 in blocking) for 1 hour at RT and mounted using ProLong Gold Antifade Mounting Media with DAPI (Invitrogen). When used, phalloidin-rhodamine (Invitrogen;R415) was incubated with secondary antibody at 1:300 concentration. Primary antibodies used were: CK7 (Millipore Sigma;CBL-194F), CK19 (Dako;M0888), ZO-1 (Invitrogen;61-7300), Cytochrome C (BD Biosciences;558709), LAMP-1 (Abcam;ab24170), KI-67 (Biolegend;350502), Acetylated a-tubulin (Millipore Signa;T6793), Gamma tubulin (GeneTex;GTX11316), Beta-catenin (GeneTex;GTX101435), CFTR (R&D Systems;MAB25031), GFP (Invitrogen;CAB4211), p62 (Abcam;ab10912), LC3 (MBL;PM036). Secondary antibodies used were Anti-Mouse Alexa 488 (Invitrogen; A32723), Anti-Mouse Alexa 647 (Invitrogen;A32728), Anti-Rabbit Alexa 594(Invitrogen;A32740), and Anti-Rabbit Alexa 555 (Invitrogen;A21428).

Organoids were grown in 25 μL droplets in 8-well Lab-Tek Chambered Coverglass (Nunc) and fixed for 20 min in 2% paraformaldehyde with 0.1% glutaraldehyde (Electron Microscopy Sciences) to prevent Matrigel depolymerization. Subsequent steps were the same as above, except longer PBS washes (3 x 45 min) were performed between each step, and organoids were counterstained with DAPI (Invitrogen) for 10 min after secondary incubation. Chambers were filled with VECTASHEILD Antifade Mounting Medium (Vector Laboratories) for imaging. Apical-out organoids were stained in 96-well plates and transferred to chamber slides in PBS and 1% low-melt agarose for imaging. All images were captured using a Leica TCS SP8 or Leica Stellaris 8 confocal microscope.

### Flow cytometry

Shortly after the completion of differentiation, iCCs were detached from the plate using Collagenase Type 2 and incubated in warm TrypLE (Gibco) to further dissociate the cells. FACS buffer (2% FBS, 1 mM EDTA, 0.1% NaN_3_) was used for all wash steps and antibody dilutions. Cells were incubated in Human TruStain FcX (Biolegend) on ice for 10 min and stained with PE Anti-Human CD326 (EpCAM) Antibody (Biolegend, #324205) at a concentration of 1:100 for 20 min on ice. DAPI was used as a marker of cell viability. Flow cytometry was performed using the BD LSR II instrument, and FlowJo software was used for analysis.

### KIF12 Gene Expression in Human Cholangiocytes

Single cell RNA sequencing data from previously published human single cell liver atlas^12^ was used to generate KIF12 expression plots across liver cell types. Single cell RNA sequencing data for primary human cholangiocytes isolated from intrahepatic bile duct, common bile duct, or gallbladder was obtained from a published study(16) (ArrayExpress accession number E-MTAB-8495). Genes identified in at least 3 cells within the dataset, cells with a minimum of 200 genes detected, and cells with <10% mitochondrial transcript percentages were included for analysis. Cells from each sample were integrated, and cholangiocytes were identified as any cell with normalized expression of EPCAM, CK7, or CK19 > 1. Seurat v. 4.1.3 was used for analysis(39).

### Transwell Cilia analysis

Images of cilia were obtained by immunostaining against Arl13b and acetylated alpha-tubulin on polarized iCCs,and were analyzed for cilia number and position. 3D reconstructions were performed on Leica LASX and Imaris (v.10.1). Cells were imaged at both low and high magnification. Cilia number and bent position were counted manually using low magnification images.

### IN/OUT Cilia analysis

The IN/OUT assay was performed as described by Kukic et. al(28). Briefly, iCCs were transduced with a lentivirus encoding an overexpression of Smoothened fused to GFP on a human PGK promoter (gifted by Toomre Lab). Cells were fixed and blocked, then incubated in anti-GFP primary antibody (1:100) for one hour. Samples were incubated in secondary antibodies (1:1000) for 1.5 hours and then mounted using Prolong Gold with DAPI. Images were analyzed using ImageJ and the plugin Cell Counter.

### Mitochondria and lysosome analysis

For immunofluorescence imaging, iCCs were plated on collagen-coated coverslips and grown to confluence as described above. Cells were stained using the above protocol with anti-LAMP1 (for lysosomes) and anti-cytochrome C (for mitochondria). For endogenous reporter imaging, lentivirus was produced using pLJC5-LAMP1-RFP-3xHA (Addgene #102932) for lysosomes and pLV-mitoDsRed (Addgene, #44386) for mitochondria. Cells were imaged 48 hours post-lentiviral transduction. Lysosome analysis was performed using an ImageJ script from Arotcarena et al.(40), used to identify the distance of each individual lysosomal puncta from the nucleus. Mitochondria analysis was performed by calculating the mitochondrial footprint using the ImageJ plugin Mitochondria Analyzer(41).

### Plasmid Construction

KIF12 coding sequence plasmid was acquired from Genecopoeia (GC-T3540). The Arg219* mutation was inserted using the N.E.B. Q5 Site Directed Mutagenesis Kit. N-terminus EGFP tagged wildtype and p.Arg219* KIF12 lentivirus vectors were constructed using Gateway Cloning LR reaction and a destination vector of pLVpuro-CMV-N-EGFP (Addgene #122848). HaloTag was constructed using restriction digest to ligate HaloTag entry vector (Addgene #107393) and KIF12 coding sequence. Gateway Cloning LR reaction was then used to insert HaloTag-KIF12 fused product in lentiviral destination vector plenti-UBC-gate-3xHA-pGK-PUR (Addgene #107393). Titration of lentivirus was calculated by tittering on GFP in iCCs aiming for 30% transfection using pRosetta plasmid. Additionally, puromycin dose was selected after 7-day trial of puromycin dose-response curve. iCCs were infected with lentivirus in the presence of polybrene (Milipore) and selected with 5μg/mL puromycin for 7 days.

### Protein expression and Western Blot

HEK293 cells were transfected with plasmids encoding N-terminal EGFP-tagged KIF12-WT and Arg219* and selected with 5 µg/mL puromycin for 7 days. The cells were harvested by centrifugation at 600g, 5 mins at 4 °C and resuspend in lysis buffer (50mM HEPES, 375mM NaCl, 20% Glycerol, Halt™ Protease Inhibitor Cocktail (Thermo Fisher Scientific), pH 7.5). Following sonication, the lysate was centrifuged for 10 mins at 17,000g, 4°C and the supernatant was collected, electrophoresed in 4–20% TGX Stain-free gel (Bio-Rad, #4568094), and transferred to a NC membrane using the iBlot 2 system (Invitrogen). Membranes were blocked in 5% non-fat milk powder solution in Tris-buffered saline and Tween-20 (TBS-T) for 1 hour at room temperature followed by overnight incubation in primary antibodies, GFP Monoclonal Antibody (Thermo Fisher) and GAPDH (Cell Signaling) at 1:1000 dilution in blocking solution 4°C. After three washes in TBS-T, the membranes were incubated with secondary antibodies, Anti-mouse IgG (NA931V) and Anti-rabbit IgG HRP (Invitrogen, #A27036) at 1:4000 dilution in blocking solution. The membrane was washed 3 times for 15 min in TBS-T and protein bands were detected with chemiluminescent substrate (Thermo Fisher, #34580) using the iBright FL 1500.

### HaloTag Single Molecule Imaging and Analysis

Serial sonication of 35mm glass-bottom dishes was performed prior to coating. The dishes were first sonicated in 1M KOH, followed by 10 minutes in MQ water and 10 minutes in 70% ethanol. After UV exposure for 20 minutes to overnight, the dishes were rinsed with PBS, air dried and then coated with filtered 50 µg/mL Collagen I. Cells were then passaged onto cleaned coated plates. Janelia Fluor® HaloTag® Ligand JFX554 (Promega) was incubated with cells at a concentration of 2nM for 15 minutes at 37°C. Cells were washed three times with PBS and returned to incubator in fresh EM media overnight. Cells were then incubated with 1:1000 dilution of Tubulin Tracker™ Deep Red (Invitrogen) for 30 minutes at 37°C, washed three times with HEPES buffer and then imaged in HEPES buffer. TIRF microscopy was performed on an Andor Dragonfly 500 spinning disk confocal microscope. B-TIRF acquisition was performed over 60 seconds at frame rate of 145.99 ms. Max projections and single molecule tracks were generated using ImageJ and plugin TrackMate(42). Kymographs were created using ImageJ to select microtubule tracks and then generated using KymographBuilder (https://imagej.net/plugins/kymograph-builder). Kymographs were analyzed using KymoButler(43) (Detection sensitivity of 0.1). Trackmate was used to generated single-molecule tracks that were further analyzed using Spot-ON kinetic modeling(27). Colocalization analyses were also performed on ImageJ using the Coloc2 (https://imagej.net/plugins/coloc-2) plugin to perform Manders’ colocalization and Costes significance tests.

### Seahorse Mitochondrial Flux Assay

Cell Mitochondrial Stress test was conducted using a Seahorse XFp analyzer (Agilent, USA) in accordance with the manufacturer’s protocol. Twenty-four hours before the assay, XFp cartridges were hydrated with XF Calibrant and incubated overnight at 37°C in a non-CO_2_ incubator. iCCs were seeded in collagen I-coated 96-well plates (Agilent). Two hours before the assay, the cells were washed three times, and the media was replaced with DMEM (Sigma) supplemented with 17.5 mM glucose, 2 mM L-glutamine, 0.2% BSA, and 10 mM HEPES (pH 7.4), followed by incubation in a non-CO_2_ incubator at 37°C. During the assay, the inhibitors were sequentially injected: 1 µM oligomycin, 10 µM FCCP, and a combination of 10 µM rotenone with 5 µM antimycin A. All inhibitors were diluted in the same DMEM medium without BSA. Hoechst 33342 dye was used to stain cell nuclei, and data were normalized per 1,000 cells using the Cyt5 integrated imaging platform (Agilent) and analyzed according to manufacturer’s protocol.

### Statistical analysis

The data represent results from at least three independent differentiations, unless indicated otherwise. The number of replicates (N) represents the data for each experiment and/or individual differentiation. Statistical analysis was performed using an unpaired t-test for all comparisons, except for Figure 3B,F, 5C and 6C, where a Mann-Whitney test was applied, and Figure 3D, which were analyzed using Welch’s test. For lysosome punctae analysis, the Kolmogorov-Smirnov test was also used. The data distribution was assessed for normality, and if both groups being compared were found to be significant, the statistical analysis was conducted accordingly. All statistical analyses were conducted with GraphPad Prism version 10.3.1, unless indicated otherwise.

