## Supplementary material for "Liver Disease Reveals KIF12 as a Critical Regulator of Mitochondria, Lysosome and Cilia Localization": file:///Users/silvlarnho/Desktop/Vilarinho/Manuscripts/KIF12-ms-II/bioRxiv/Supplementary-Information-2-24-2026

**Supplementary Material and Methods**

***Bulk RNA-seq and analysis***

RNA was isolated from cells at hepatic endoderm and cholangiocyte-like stages using the RNeasy Plus Mini Kit (Qiagen). RNA was submitted to the Yale Center for Genome Analysis for bulk RNA sequencing with ribosomal reduction using the Illumina Novaseq platform (25M reads per sample). Raw reads were trimmed using Trimmomatic v.0.39 and then aligned to the GRCh38 human reference genome using STAR (v. 2.7.7a). Gene counts were generated using HTSeq (v. 0.13.5), and DESeq2 package^39^ in R was used for differential expression analysis with variance stabilizing transformation. Pairwise comparisons between conditions were performed using a Wald test, while comparisons between all samples were performed using a likelihood-ratio test. Principal component analysis (PCA) visualization was performed using the *pca* function from the M3C package^40^ in R. Differentially expressed genes were identified as genes with Benjamini-Hochberg-corrected adjusted p-value < 0.05 and log2 fold-change (logFC) > 1. Gene ontology enrichment analyses were performed using the clusterProfiler package^42^ in R. Gene set enrichment analysis (GSEA) was performed using the genekitr package^43^ in R. To determine similarity with published RNA-Seq, ARCHS4 was used to pull data from GEO for GSE86007, GSE120732, GSE146899, GSE157791 and GSE182877. Data was then normalized using ComBat within the sva package. Gene scores were calculated by using the log counts per million mean score of the following genes for each subset: intrahepatic cholangiocytes (*SOX4*, *DCDC2*, *PKHD1*, *HES4*, *BICC1*, *JAG1*, *ANKRD1*) and extrahepatic cholangiocytes (*SOX17*, *AQP1*, *FGF19*, *SLC15A1*, *MUC13*, *SLC12A7*)^14^.

***Secretin swelling assay***

Two days prior to performing the secretin assay, forskolin was removed from EM media to minimize forskolin-induced swelling of the organoids. Brightfield images of each organoid were taken at 15 min intervals using a 20X objective before and after the addition of 100 nM Secretin (Sigma). Organoids were incubated at 37°C between each time point. ImageJ was used to determine the percent increase in lumen area.

***Autophagy analysis***

For p62 immunofluorescent imaging, iCCs were plated on collagen-coated coverslips and grown to confluence as described above. Cells were stained using the above protocol with anti-p62. ImageJ “Find Maxima” function was used to count number of p62 positive of repunctae. For DQ-BSA analysis, cells were similarly plated on collagen-coated coverslips and grown to confluence. Cells were incubated with 10mg/mL DQ-BSA (Thermo Fisher, #D12051) for 2 hours at 37°C and then washed twice with PBS. Cells were fixed with 4% PFA and then imaged using immunofluorescence as previously described. ImageJ was used to calculate relative fluorescence. For LC3 immunofluorescent imaging, iCCs were plated on collagen-coated coverslips and grown to confluence. Cells were incubated with either DMSO or 100nM bafilomycin (Millipore-Sigma #19-148) for 1 hour, subsequently washed and then fixed. Cells were stained using the above protocol with anti-LC3. ImageJ “Find Maxima” function was used to count number of p62 positive punctae.

**Supplementary Figures Legends**

**Figure S1.** **Expression of 45 human kinesins across liver cell types using aggregated and integrated scRNA-seq data from 28 non-diseased human livers.**

**Figure S2.** **Characterization of Y6 wildtype and Arg219* KIF12 biliary organoids.** (A) Representative images of CK7 and CK19 immunostaining of biliary organoids. (B) Quantification of the average change in lumen area of wildtype and Arg219* biliary organoids upon incubation with secretin. Photomicrographs depict representative organoid swelling after 120 minutes. n > 18 organoids per genotype from two independent differentiations.

**Figure S3.** **Transcriptomic analysis of Y6 wildtype and *KIF12* mutant Arg219* iPSC-derived hepatic endoderm (iHE) and cholangiocyte-like cells (iCCs).** (A) Principal component analysis (PCA) of bulk RNA sequencing of Y6 wildtype and KIF12 Arg219* iHEs (3 independent differentiations) and iCCs (2 independent differentiations). (B) PCA plot of RNA sequencing of Y6 wildtype and KIF12 Arg219* iCCs, previously published iCCs from another group^23^, human primary cholangiocytes, and the NHC cell line. (C) Normalized gene score of intra-hepatic cholangiocytes (IHC) representative genes and extra-hepatic cholangiocytes (EHC) genes within Y6 WT and KIF12 Arg219* iCCs. (D,E,F) Gene ontology enrichment analyses of DEGs between Y6 wildtype and KIF12 Arg219* iCCs for cellular components, biological processes, and molecular functions, respectively. Circle color represents the significance of the pathway, while circle size denotes the number of DEGs enriched within the pathway.

**Figure S4. TIRF microscopy of single KIF12 wildtype molecules in Y6 iCCs.** (A) Representative images of TIRF microscopy of Tubulin Tracker highlighting microtubules (grey) and wildtype Halo-KIF12 overexpression (magenta) in live iCCs. Single frames (left panel) and max projection from 1-minute acquisition (right panel) are shown. (B) Additional representative kymographs of wildtype Halo-KIF12 along a single microtubule. Left kymograph depicts Halo-KIF12, middle kymograph depicts Tubulin Tracker highlighting microtubules, right kymograph is a composite image.

**Figure S5. Mitochondrial location in Y6 *KIF12* Arg219* iCCs.** Overexpression of MitoDsRed in iCCs illustrating mitochondrial perinuclear localization and footprint reduction in Arg219* iCCs. Corresponding bar graphs show the quantified mitochondrial footprint, with each dot representing the average percentage per experiment. Data are presented as mean ± S.D (n=3-4).

**Figure S6. Overview of second independent FL167 iPSC *KIF12* Arg219* line.** (A) Schematic representation of CRISPR strategy for introduction of homozygous Arg219* mutation in *KIF12* in FL167 iPSC line. (B) Sanger sequencing of wildtype FL167 (top) and KIF12 Arg219* FL167/F11 (bottom) iPSC lines demonstrating homozygous C>T mutation in the latter. (C) Surface expression of epithelial marker EpCAM in FL167-derived iCCs. (D) Representative images of CK7 (green), ZO-1 (red), and phalloidin (yellow) staining in KIF12 wildtype (FL167) and Arg219* (FL167/clone-F11) iCCs.

**Figure S7.** **Lysosomes in Y6 *KIF12* Arg219* iCCs.** (A) Representative image of LAMP1-RFP overexpression in iCCs. Violins represent the distribution of the distance of individual LAMP1^+^ puncta from the nucleus. Quantification includes >1450 LAMP1-RFP puncta per genotype (n≥3). ****p≤0.0001. (B) Representative images of p62 immunostaining (red) and nucleus (blue). Each dot represents quantification of p62 punctae per one image, including > 100 cells per genotype from two independent experiments. (C) Representative images of DQä-BSA uptake in autophagosomes. Each dot represents relative fluorescence per image, including >250 cells per genotype from two independent experiments. (D) Representative images of LC3 immunostaining (red) and nucleus (blue) after treatment with DMSO or 100nM of bafilomycin. Each dot represents LC3 positive punctae per image, including >250 cells per genotype from two independent experiments.

**Figure S8.** **Primary cilia are bent in Y6 *KIF12* Arg219* iCCs.** (A) Representative Z-stack images of ARL13β (red) immunostaining highlighting cilia in iCCs. Separate higher magnification images of the selected views display two cells with cilia in two distinct orientations: upright (top panel) and bent (lower panel). (B) Quantification of percentage of bent cilia using ARL13β immunostaining with >1200 cilia per genotype.

**Figure S9.** **Apical-Out Organoids Derived from Y6 wildtype and Arg219* KIF12 iCCs.** (A) Representative images of 'apical-out' (inverted) biliary organoids displaying reversed phalloidin (cyan) along with acetylated α-tubulin (green) and ARL13β (red), illustrating cilia misorientation in Arg219* mutants and apical location in wildtype controls. (B) Images of Apical-out biliary organoids stained for CFTR (green), β-catenin (red), and phalloidin (cyan), demonstrating intact membrane polarity in both wildtype and Arg219* KIF12 organoids.

**Figure S10.** **Primary cilia in FL167 wildtype and Arg219* KIF12 derived biliary organoids.** (A) Biliary organoids stained with phalloidin (green) and β-catenin (red) illustrate intact apical-basal polarity in both FL167-derived wildtype and Arg219* iCCs. Co-immunostaining with ciliary markers acetylated α-tubulin (green) and ARL13β (red) reveals apical cilia in wildtype biliary organoids and basolateral cilia in KIF12 mutant biliary organoids (B) 'Apical-out' (inverted) biliary organoids showing reversal of phalloidin (green) and β-catenin (red) staining, along with acetylated α-tubulin (green) and ARL13β (red) co-localization on both the apical and basal sides, highlighting cilia misorientation in Arg219* despite maintaining normal apical-basal polarity. White arrows indicate cilia. Scale bars= 20 microns.

**Figure S11.** **Wildtype KIF12 overexpression restores bent cilia in Y6 Arg219* iCCs.** (A) Representative Z-stack images of overexpression (OE) of EGFP-tagged Arg219* and wildtype KIF12 constructs in Y6 iCCs immunostained with ARL13β (red), showing restoration of cilia to the upright position in Arg219*-Wildtype OE iCCs. Separate higher magnification images of the selected views display cilia in two distinct orientations: bent (top panel) and upright (lower panel). (B) Quantification of the proportion of bent cilia using ARL13β immunostaining with >580 cilia per genotype, (n=5). ****p≤0.0001. Error bars represent mean ± S.D

**Supplementary Videos**

**Video S1**. 3D reconstruction of Z-stack images of ARL13β (red) immunostaining highlighting cilia and DAPI (blue) in wildtype iCCs.

**Video S2**. 3D reconstruction of Z-stack images of ARL13β (red) immunostaining highlighting cilia and DAPI (blue) in Arg219* iCCs.

**Video S3**. 3D reconstruction of Z-stack images of ARL13β (red) immunostaining highlighting cilia and DAPI (blue) in Arg219* iCCs rescued with wildtype KIF12 overexpression.

**Video S4**. 3D reconstruction of Z-stack images of ARL13β (red) immunostaining highlighting cilia and DAPI (blue) in Arg219* iCCs with Arg219* mutant KIF12 overexpression.
