## Supplementary figures and images for "Liver Disease Reveals KIF12 as a Critical Regulator of Mitochondria, Lysosome and Cilia Localization"

### file:///Users/silvlarnho/Desktop/Vilarinho/Manuscripts/KIF12-ms-II/bioRxiv/Supplementary-Figures/PDFs/Supplementary-Figure-1.pdf

Supplementary Figure 1

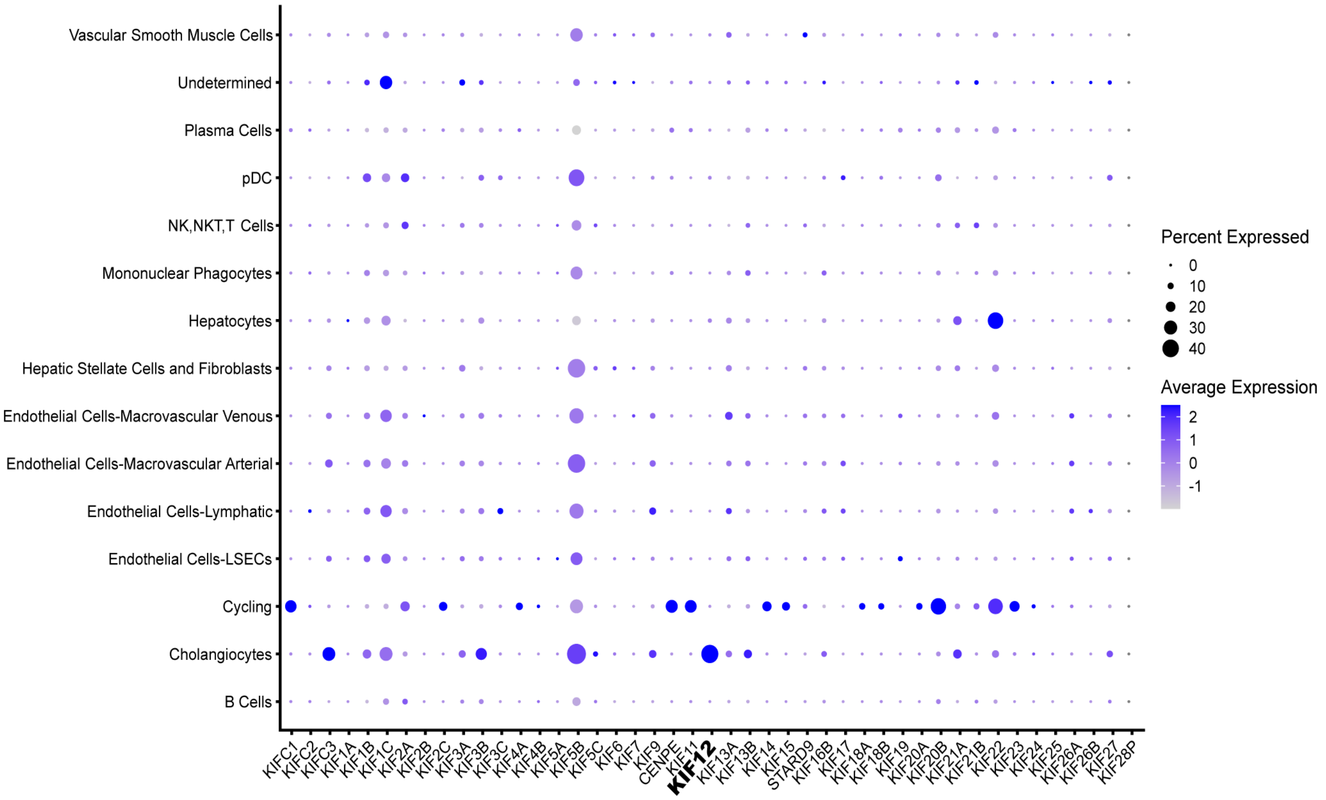

### file:///Users/silvlarnho/Desktop/Vilarinho/Manuscripts/KIF12-ms-II/bioRxiv/Supplementary-Figures/PDFs/Supplementary-Figure-2.pdf

Supplementary Figure 2

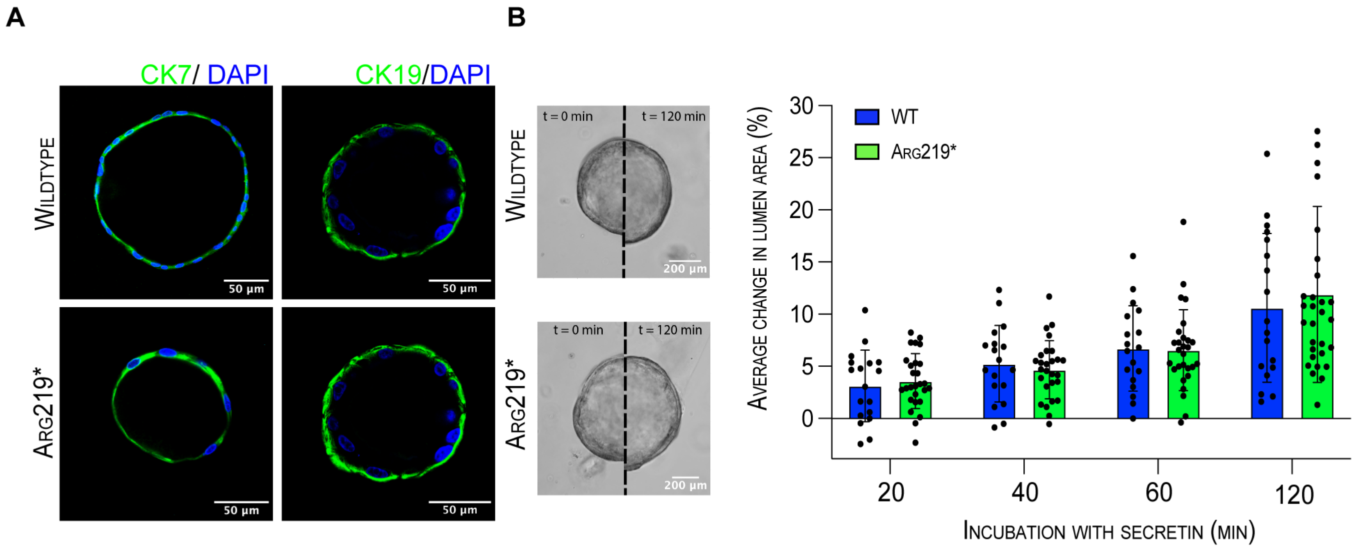

### file:///Users/silvlarnho/Desktop/Vilarinho/Manuscripts/KIF12-ms-II/bioRxiv/Supplementary-Figures/PDFs/Supplementary-Figure-3.pdf

#### Supplementary Figure 3

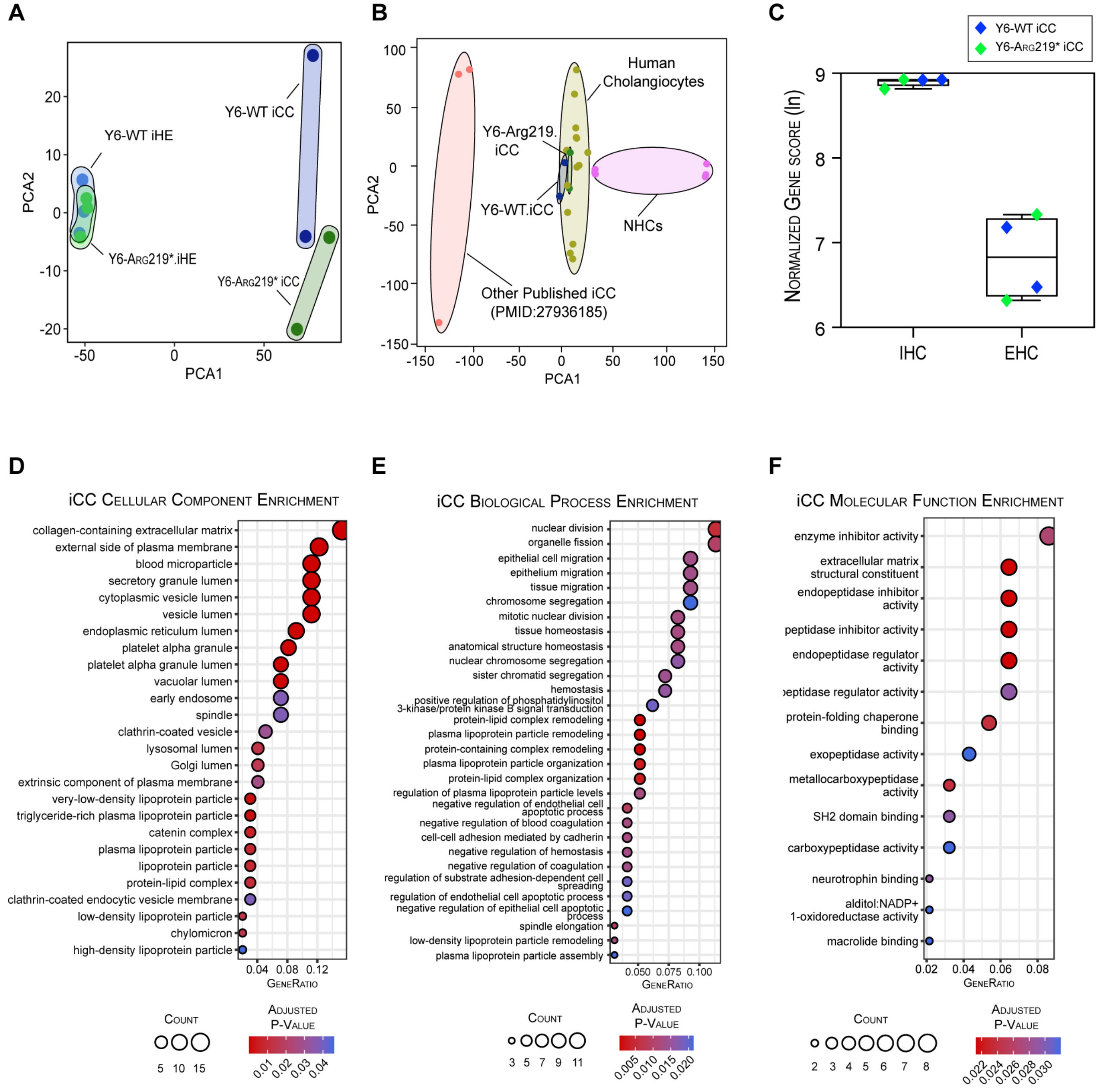

### file:///Users/silvlarnho/Desktop/Vilarinho/Manuscripts/KIF12-ms-II/bioRxiv/Supplementary-Figures/PDFs/Supplementary-Figure-4.pdf

Supplementary Figure 4

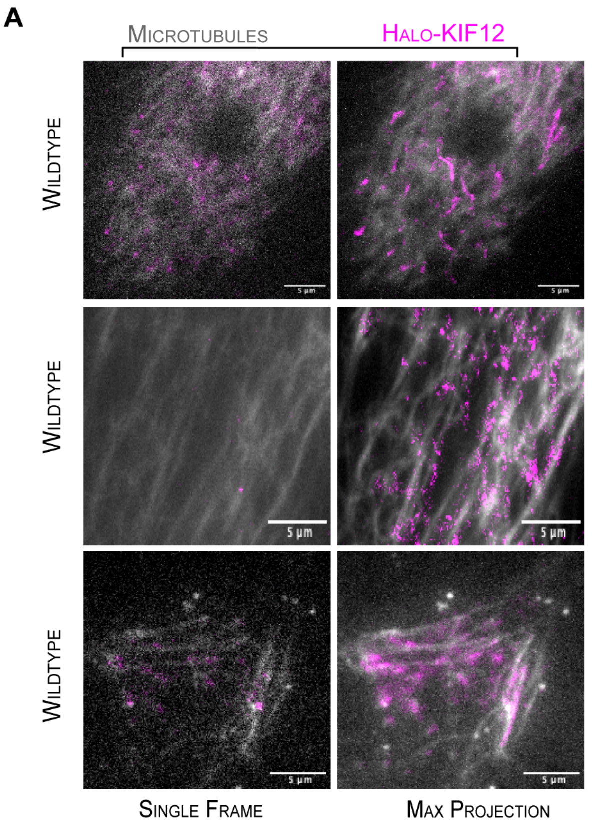


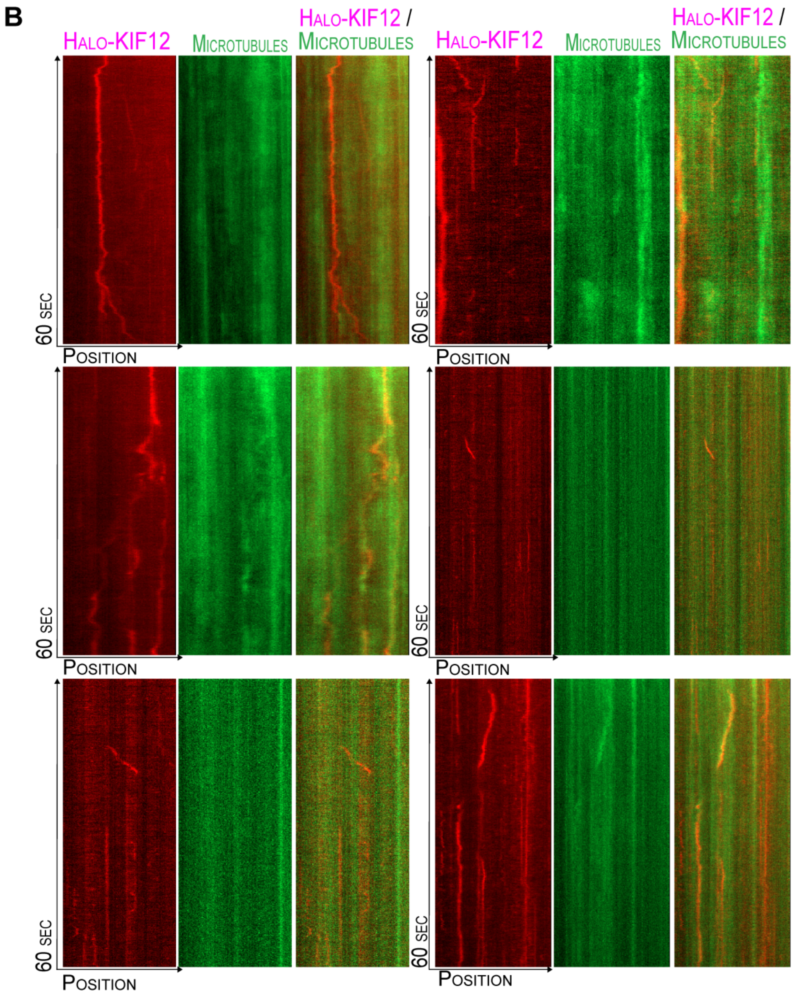

### file:///Users/silvlarnho/Desktop/Vilarinho/Manuscripts/KIF12-ms-II/bioRxiv/Supplementary-Figures/PDFs/Supplementary-Figure-5.pdf

Supplementary Figure 5

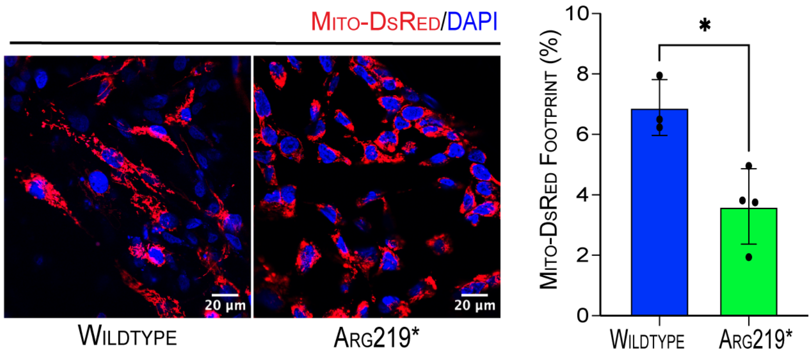

### file:///Users/silvlarnho/Desktop/Vilarinho/Manuscripts/KIF12-ms-II/bioRxiv/Supplementary-Figures/PDFs/Supplementary-Figure-6.pdf

Supplementary Figure 6

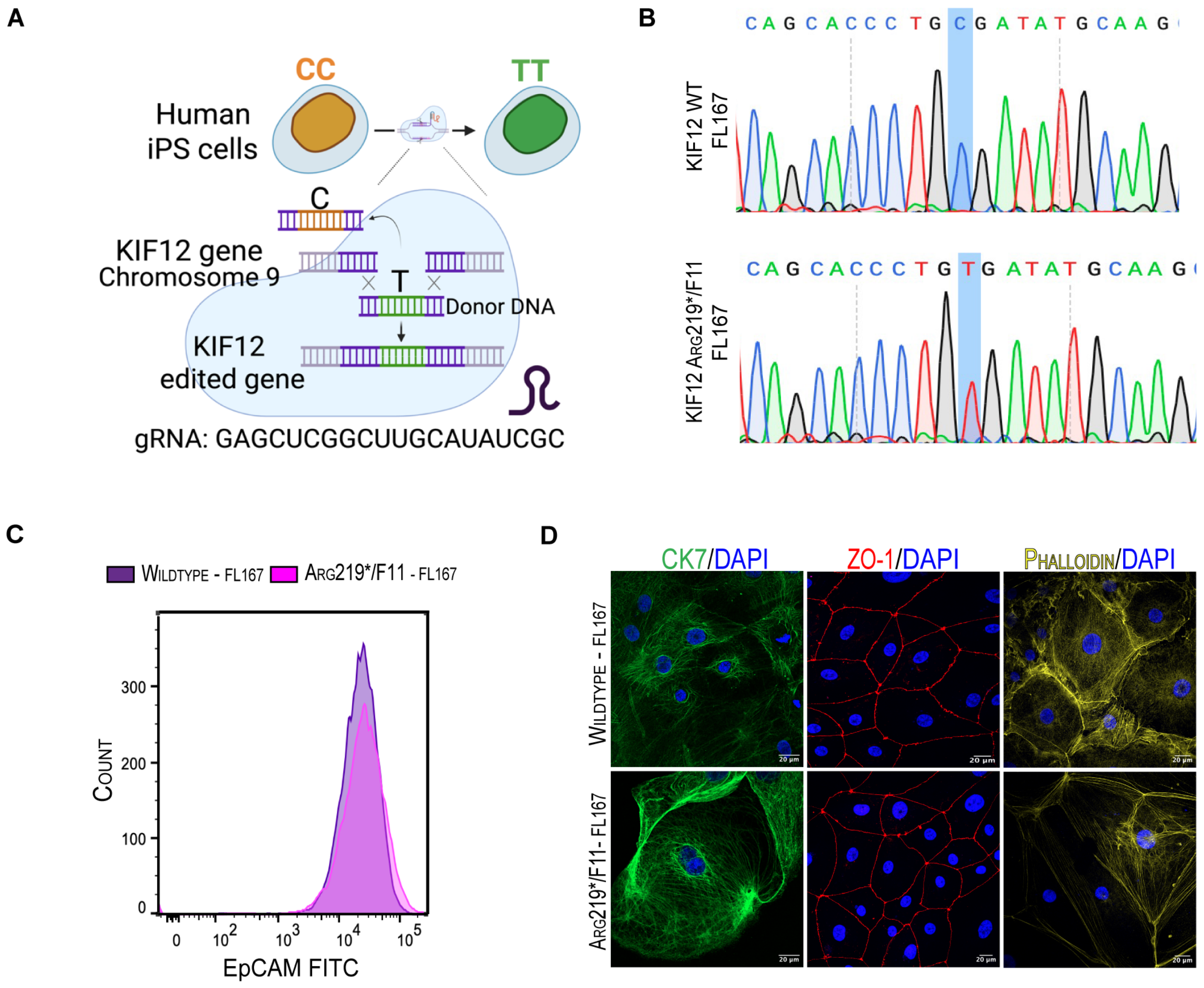

### file:///Users/silvlarnho/Desktop/Vilarinho/Manuscripts/KIF12-ms-II/bioRxiv/Supplementary-Figures/PDFs/Supplementary-Figure-7.pdf

Supplementary Figure 7

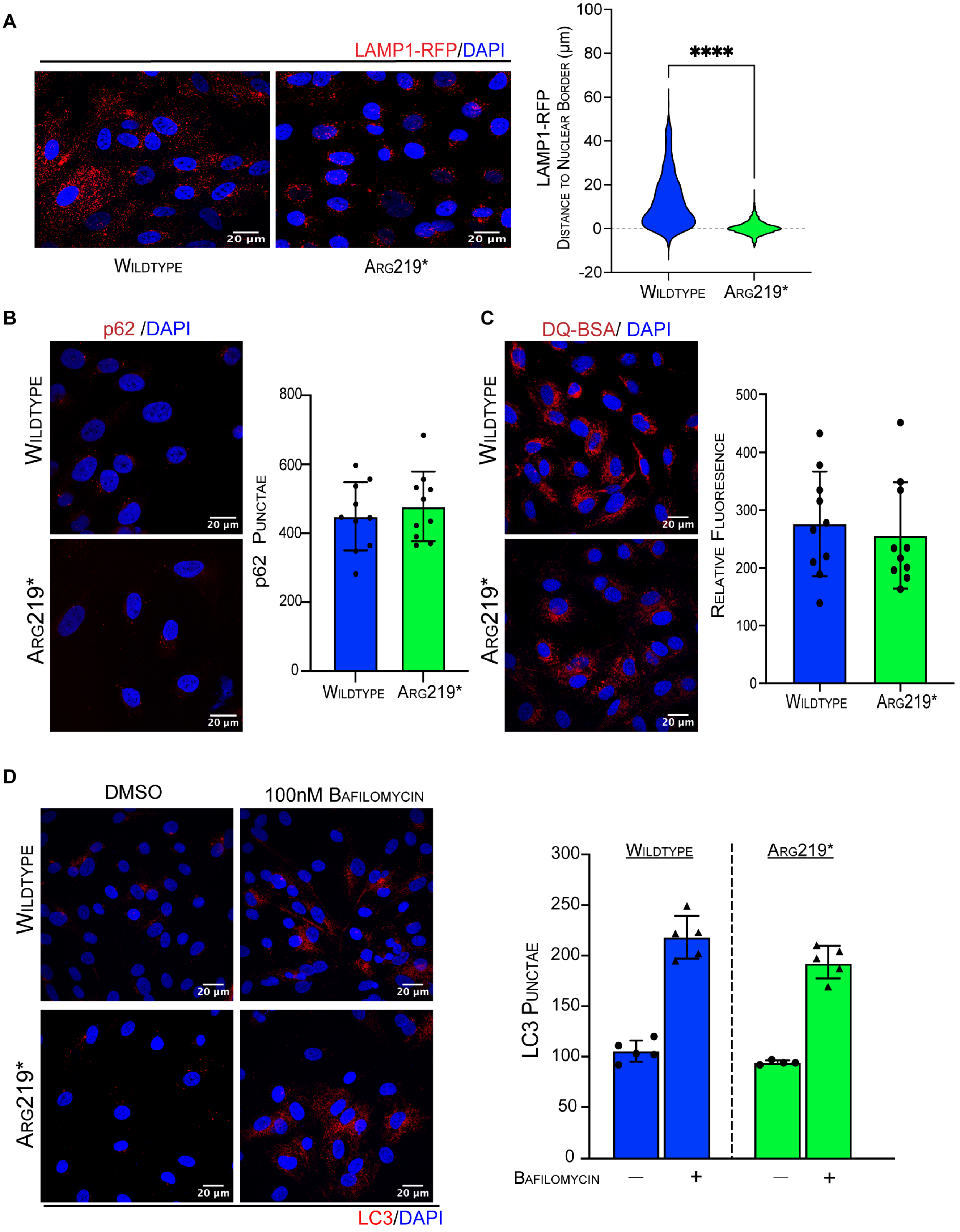

### file:///Users/silvlarnho/Desktop/Vilarinho/Manuscripts/KIF12-ms-II/bioRxiv/Supplementary-Figures/PDFs/Supplementary-Figure-8.pdf

Supplementary Figure 8

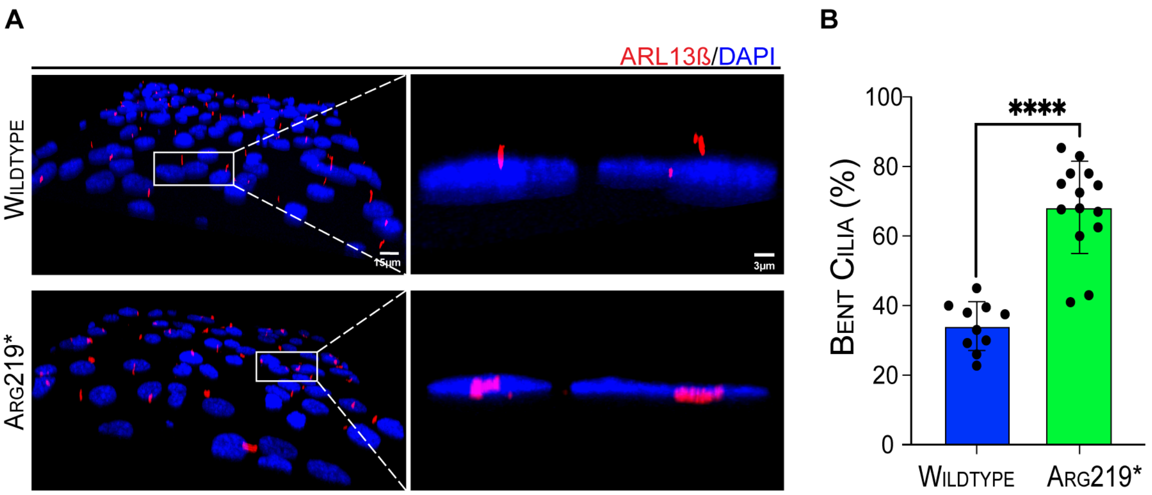

### file:///Users/silvlarnho/Desktop/Vilarinho/Manuscripts/KIF12-ms-II/bioRxiv/Supplementary-Figures/PDFs/Supplementary-Figure-9.pdf

Supplementary Figure 9

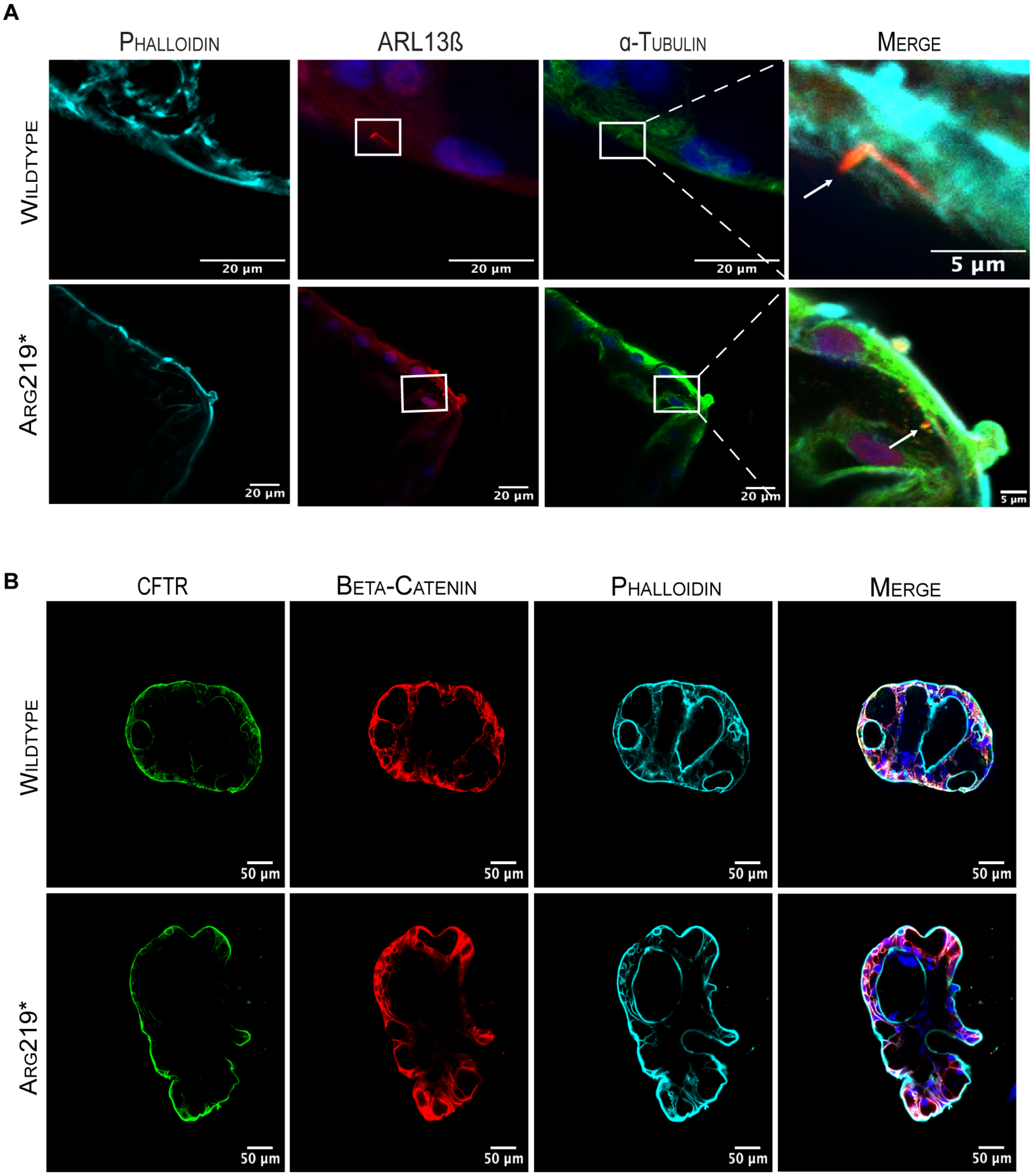

### file:///Users/silvlarnho/Desktop/Vilarinho/Manuscripts/KIF12-ms-II/bioRxiv/Supplementary-Figures/PDFs/Supplementary-Figure-10.pdf

Supplementary Figure 10

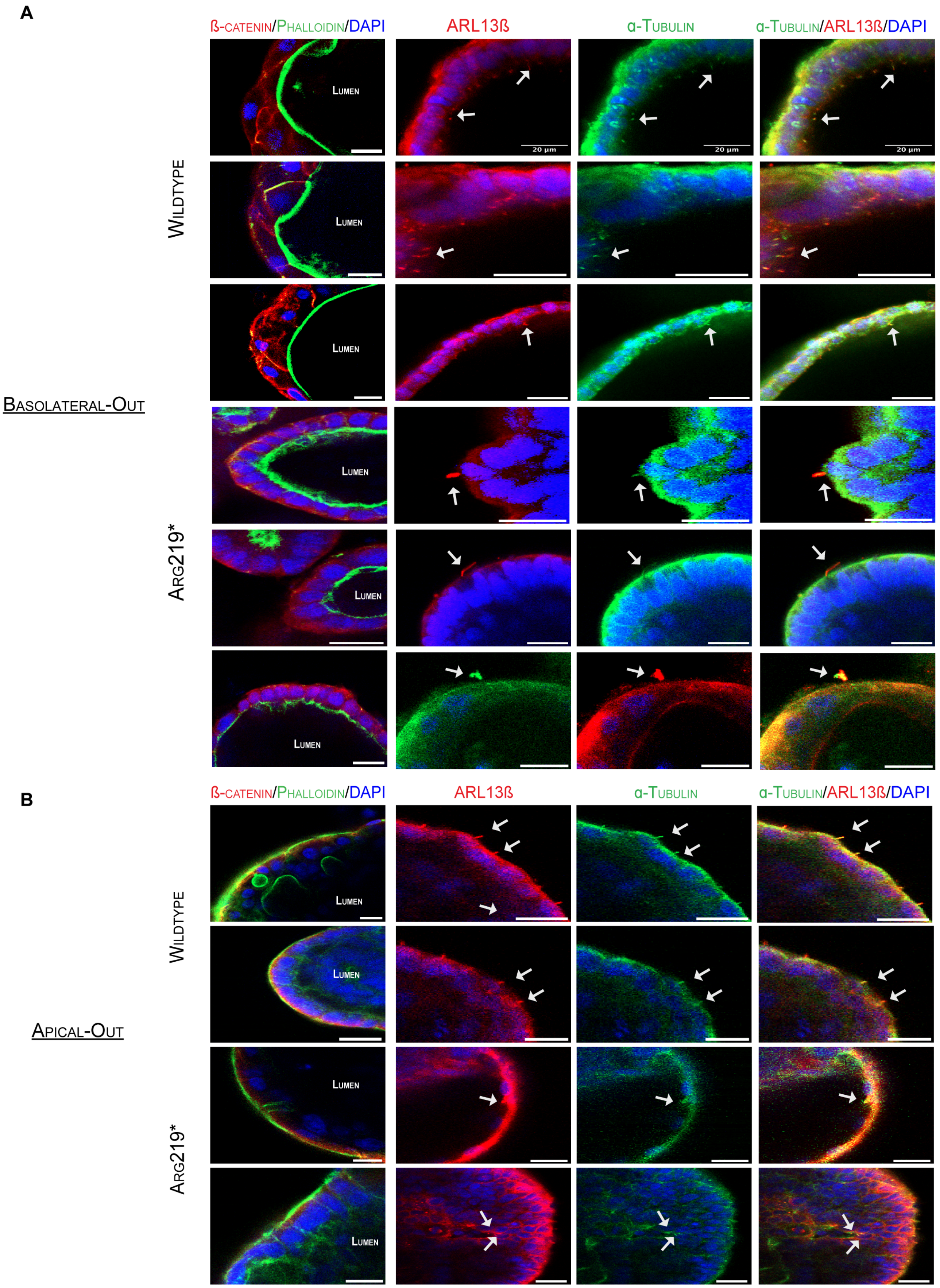

### file:///Users/silvlarnho/Desktop/Vilarinho/Manuscripts/KIF12-ms-II/bioRxiv/Supplementary-Figures/PDFs/Supplementary-Figure-11.pdf

Supplementary Figure 11

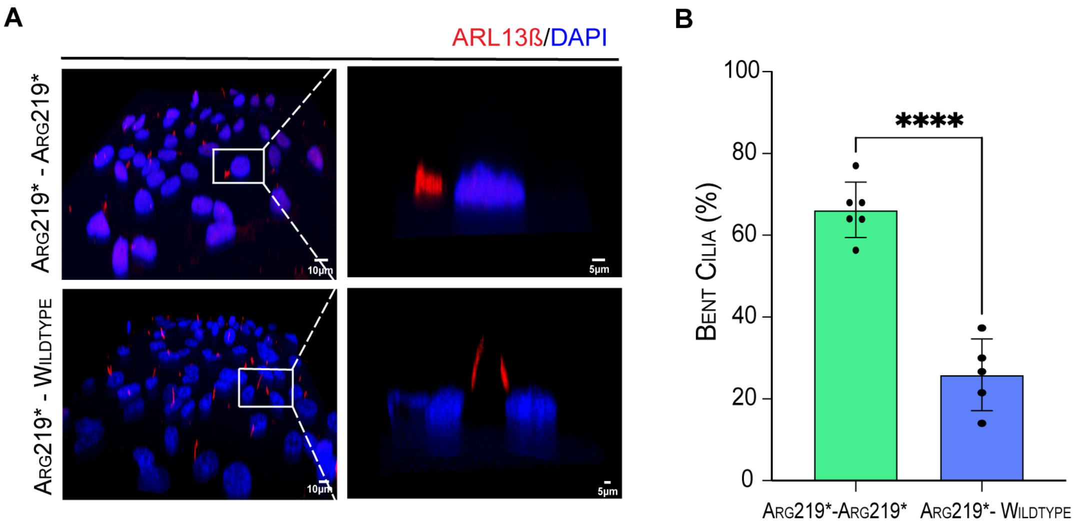
